# Identifying non-identical-by-descent rare variants in population-scale whole genome sequencing data

**DOI:** 10.1101/2020.05.26.117358

**Authors:** Kelsey E. Johnson, Benjamin F. Voight

## Abstract

The site frequency spectrum in human populations is not accurately modeled by an infinite sites model, which assumes that all mutations are unique. Despite the pervasiveness of recurrent mutations, we lack computational methods to identify these events at specific sites in population sequencing data. Rare alleles that are identical-by-descent (IBD) are expected to segregate on a long, shared haplotype background that descends from a common ancestor. However, alleles introduced by recurrent mutation or by non-crossover gene conversions are identical-by-state and will have a shorter expected shared haplotype background. We hypothesized that the expected difference in shared haplotype background length can distinguish IBD and non-IBD variants in population sequencing data without pedigree information. We implemented a Bayesian hierarchical model and used Gibbs sampling to estimate the posterior probability of IBD state for rare variants, using simulations to demonstrate that our approach accurately distinguishes rare IBD and non-IBD variants. Applying our method to whole genome sequencing data from 3,621 individuals in the UK10K consortium, we found that non-IBD variants correlated with higher local mutation rates and genomic features like replication timing. Using a heuristic to categorize non-IBD variants as gene conversions or recurrent mutations, we found that potential gene conversions had expected properties such as enriched local GC content. By identifying recurrent mutations, we can better understand the spectrum of recent mutations in human populations, a source of genetic variation driving evolution and a key factor in understanding recent demographic history.

## Introduction

Recurrent mutations are repeated mutational events at the same nucleotide position in multiple individuals in a population. The frequency of recurrent mutations and their relevance to evolutionary genetics studies have been examined since the beginning of the field of population genetics (e.g. Wright 1931; Haldane 1933; Wright 1937). The frequency that a recurrent mutation is observed in a sample depends on many factors, including the per-base-pair mutation rate, the number of chromosomes surveyed, the effective population size, as well as the demographic history of the population surveyed. Distinguishing recurrent mutations from variants whose alleles are all inherited identical-by-descent (IBD) is critical to a complete understanding of the human germline mutation rate, and to population genetic methods that make inferences from the observed number and frequency of genetic variants in a population.

As the genetics community has surveyed large, rapidly growing populations with a finite genome size, such as modern humans (Harpak et al. 2016), it has been observed that recurrent mutations occur at appreciable frequency. For example, in the Exome Aggregation Consortium dataset of 60,706 human exomes, there is a marked absence of singleton CpG transitions relative to other mutation types (Lek et al. 2016). This observation could be explained by the presence of recurrent mutations saturating these highly mutable sites in this large sample, resulting in two or more sampled individuals segregating identical-by-state alleles at CpG sites.

One implication of this observation is that the presence of recurrent mutations in a large sample may result in a suboptimal calibration of summary data typically utilized in population genetic inference, like the site frequency spectrum (SFS). The SFS is the distribution of the number of observed variants at allele counts 1 to *n*-1 in a sample of *n* chromosomes. Many modern population genetics methods use the SFS to infer the demographic history of a sample (Gutenkunst et al. 2009; Lukic and Hey 2012; Excoffier et al. 2013; Bhaskar et al. 2015; Jouganous et al. 2017). These approaches generally assume an infinite sites model with no recurrent mutations, but the human site frequency spectrum is not well explained by an infinite sites model (Harpak et al. 2016). Previous work has described the SFS allowing for recurrent mutations, relying on observed recurrent mutations in the form of triallelic sites (Jenkins and Song 2011; Jenkins et al. 2014; Ragsdale et al. 2016). If recurrent mutations are not accounted for, the SFS will be shifted to higher allele frequencies relative to the SFS that incorporated recurrent mutations at lower frequencies. This could impact the accuracy of demographic parameter inference. In particular, methods that infer the magnitude of recent population growth, which rely on rare variants, may incur bias if they do not take recurrent mutations into account. Similarly, the magnitude of purifying selection may be underestimated if the frequencies of rare variants are overestimated due to undetected recurrent mutation. Finally, estimates of mutation rates from population genetic data that do not incorporate recurrent mutations may be biased downwards.

Beyond population genetic applications, identifying specific recurrent mutations could be useful in the context of genotype-phenotype association through tests of rare variant burden. Rare variant burden approaches are increasingly used to associate genomic regions with disease status or quantitative traits in large-scale sequencing datasets (Nicolae 2016). In general, these approaches test the null hypothesis that the frequency of rare variants in a genomic region is independent of the phenotype of interest. If a gene is causal for a trait, we may expect to observe a higher frequency of rare variants, but also more recurrent variants especially at highly mutable nucleotide positions that have a substantial impact on the trait. If recurrent mutations could be identified, they could potentially improve power to associate the gene with the trait. Recurrent mutations have been used in this context in family-based studies, where recurrent mutations can be identified as *de novo* events in unrelated families (e.g. Kirby et al. 2013; O’Roak et al. 2014). We are not aware of any examples of recurrent mutations being used in large-scale population-based sequencing studies of rare disease associations.

In what follows, we propose a computational approach to infer the presence of a recurrent mutation at a genomic site. The key idea underlying our approach is to use the genetic variation linked to rare variants to distinguish alleles at a variant position as identical-by-descent (IBD) or non-IBD. Rare IBD variants are usually surrounded by a long, shared haplotype on all chromosomes carrying the variant (i.e., an IBD segment), because all segregating alleles derive from a recent ancestral mutation. If the variant arose a small number of generations ago, there have been few opportunities for recombination events to shorten the shared IBD segment. Thus, the length of the IBD segment shared across carriers is inversely related to the age of the variant (Haldane 1919; Mathieson and McVean 2014). In contrast, recurrent mutations or gene conversions can occur on any random haplotype background in a population, and thus we expect that their local time to the most recent common ancestor (TMRCA) will be on average older than an IBD variant of the same allele frequency.

Leveraging the relationship between local TMRCA and the length of a shared IBD segment, we can identify rare variants that appear non-IBD. However, it is important to consider many potential reasons why we might observe a short IBD segment around a rare variant. Beyond recurrent mutation, non-crossover gene conversion, proximity to a region of extremely high local recombination rate, or simply chance might also explain specific events. In addition, if one or multiple copies of a rare variant are genotyping errors, this could result in the same signature of a shorter than expected shared IBD segment between carriers. Thus, any approach that aims to identify recurrent mutations from data must work to distinguish amongst these types of events.

We propose to identify rare variants that fall on the extreme short end of shared IBD segment lengths and attempt to categorize these likely non-IBD variants by the possibilities enumerated above. In addition to the IBD segment length, additional genomic annotations can help to distinguish between these causes, such as the mutation’s sequence context (e.g. a CpG mutation), the local recombination rate, and local GC content.

While previous efforts have leveraged IBD tracts to infer mutation and gene conversion rates (Palamara et al. 2015), as well as to estimate allele ages (Palamara et al. 2012; Mathieson and McVean 2014; Platt et al. 2019; Albers and McVean 2020), we are not aware of any previous method designed to specifically identify recurrent mutations and gene conversion events at specific genomic positions at genome-wide scale. In today’s era of whole genome sequencing of thousands of individuals from a population, categorizing specific rare variants as likely recurrent mutations or gene conversions is now uniquely possible. Here, we describe a Bayesian hierarchical model to identify non-IBD rare variants, using population genetic simulations to assess its precision and accuracy. We then apply our approach to sequencing data of 3,621 individuals from the UK10K dataset, and partition high-confidence non-IBD rare variants as those likely to be recurrent mutations or gene conversions.

## New Approaches

### Theory

Previous work to model the expected TMRCA between two IBD alleles or two random alleles in a population provides a framework through which these states can be distinguished in data. Measuring the accumulation of mutations on a haplotype, *i.e.*, the mutational clock, is useful for estimating the age of older, common variants; however, for our purpose here to distinguish rare recurrent and IBD variants, there will be few if any linked mutations more recent than the focal variant. Therefore, the mutational clock does not help us distinguish IBD and non-IBD rare alleles, and we rely solely on the recombination clock for inference.

The theoretical distributions of the pairwise TMRCAs for IBD or non-IBD alleles for a range of allele counts are plotted in **Supplementary Figure 1** (**Supplementary Methods**). As the IBD allele count increases, the mean TMRCA also increases, reflecting that higher frequency alleles tend to be older; meanwhile, the TMRCA distribution between non-IBD allele pairs is unchanging because it is not a function of the allele frequency. Thus, the difference between the expected TMRCA for IBD vs. non-IBD variants increases with decreasing allele frequency.

Though the TMRCA of a genetic variant is not directly observable, it can be estimated by the length of the haplotype shared by carriers of the variant. The distance to the nearest recombination event on either side of a genetic variant between a pair of alleles can be modeled as exponentially distributed with rate proportional to the TMRCA (Palamara et al. 2012; Mathieson and McVean 2014). The expected difference in the TMRCA between a rare variant segregating with IBD alleles and a variant position with recurrent mutations (**Supplementary Figure 1**) translates into IBD variants having, on average, longer pairwise distances to obligate recombination events compared to recurrent sites of the same allele frequency (**Supplementary Figure 2**). Recent methods have inferred the age of alleles in large-scale population datasets by leveraging this relationship between haplotype background and the TMRCA (Palamara et al. 2012; Platt et al. 2019; Albers and McVean 2020), and by constructing local genealogies (Kelleher et al. 2019; Speidel et al. 2019). However, these tools assume an infinite sites model (*i.e.*, no recurrent mutations), and do not explicitly attempt to identify recurrent mutations. Existing approaches to identify recurrent mutations rely on family relationships, or assume that variants present at very rare frequencies in distantly related populations are recurrent without explicitly identifying variants as non-IBD (e.g. Pagnier et al. 1984; The 1000 Genomes Project Consortium 2012). Thus, our goal was to develop an approach to identify non-IBD variants that scales to large, whole genome population sequencing studies with thousands of individuals.

While recombination breakpoints cannot be directly observed in population sequencing data, patterns of genetic variation can give us an estimate of the location of these events. Here, we are interested in rare genetic variants that are difficult to accurately phase. Additionally, the signature of recurrent mutation itself could introduce error into statistical phasing algorithms. Thus, our method utilizes unphased diploid genotypes to estimate the recombination distances on either side of a pair of alleles. With diploid genotypes for a pair of individuals each carrying a focal allele, one can measure the obligate recombination distance as the physical span to the first opposite homozygote genotype between the two individuals (**Supplementary Figure 3**). No genealogy without recombination is compatible with the observed genotypes of these two sites (the focal allele and the site of the opposite homozygote genotypes), and so we assume a recombination event has occurred between them (Mathieson and McVean 2014). Thus, the obligate recombination distance gives an estimate of the true recombination distance.

### Statistical Model

We considered a Bayesian hierarchical model for the pairwise recombination distances from a sample of variants of a given allele count, which allowed us to learn the model parameters directly from the data (**Figure 1**). We modeled the sampled variants as a finite mixture of IBD (*k*=1) or non-IBD (*k*>1, with each possible partition of alleles for a non-IBD variant given a different value of *k*), with mixture proportions *π*_*k*_. For example, a non-IBD variant of allele count 4 has two possible partitions: a singleton and an IBD tripleton (1:3), or two doubletons (2:2). Each variant of allele count *A* has 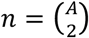 allele pairs. The TMRCA (*t*) for an allele pair was sampled from a gamma distribution with shape *α* and rate *β*, with one gamma distribution of *t* for IBD allele pairs and one for non-IBD allele pairs, For non-IBD allele pairs, we estimated *α* and *β* from multiallelic sites, and for IBD allele pairs we fixed *α* and performed sampling for *β*, over a range of possible values for *α*. For each variant, the possible permutations of allele pair assignments of IBD or non-IBD states are denoted by *j.* For IBD variants, all allele pairs are IBD; for non-IBD variants, the possibilities depend on the partition *k*. We modeled the left and right recombination distances (*d*_*L*_, *d*_*R*_) for each allele pair following an exponential distribution with rate proportional to *t*. We used Gibbs sampling to sample from the marginal posterior density of each parameter, as we could estimate these densities from the full conditional distributions. Below we outline these expressions.

**Figure 1.**
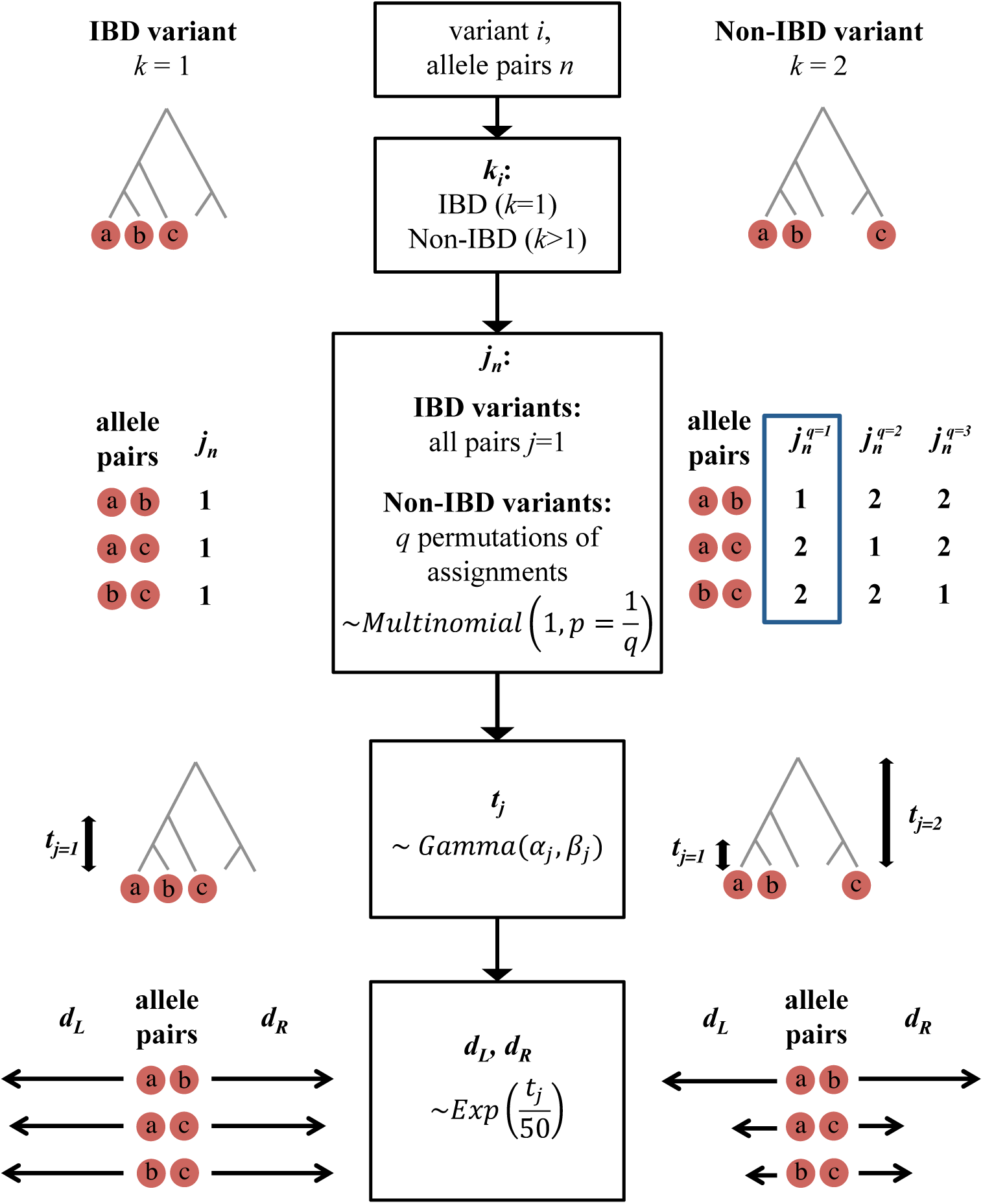
The generative model underlying our Bayesian hierarchical model to distinguish IBD and non-IBD variants. Each variant *i* has *n* allele pairs; *k*: variant assignment to IBD (k=1) or non-IBD (k>1); *j*_*n*_: allele pair assignments (IBD: *j*_*n*_ =1, non-IBD: *j*_*n*_ =2); *q*: all possible permutations of *j*_*n*_ assignments for a given non-IBD variant partition; *t*_*j*_: within a variant, IBD allele pairs or non-IBD allele pairs’ TMRCAs; *d*: allele pairwise recombination distances to the right (*d*_*R*_) and left (*d*_*L*_).

#### Mixture proportions (π)

Using a multinomial likelihood for the probability of the assignments *k* based on proportions *π*, we used the conjugate prior Dirichlet distribution to get a Dirichlet posterior for the probabilities of *π* given the observed *k* assignments. Thus we have the likelihood function:

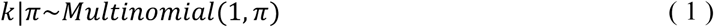

We used a Dirichlet prior for *π*:

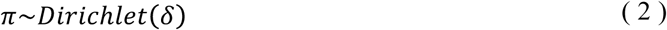

The resulting posterior probability followed a Dirichlet distribution:

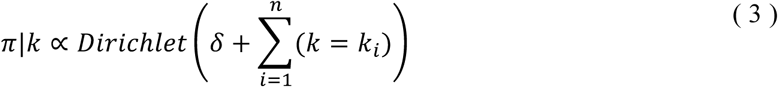

#### TMRCA (t)

The likelihood of the pairwise recombination distance *d* to one side of a variant (in centiMorgans), given TMRCA *t*, followed an exponential distribution (Palamara et al. 2012):

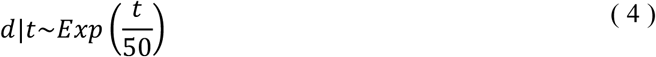

We used a gamma prior for *t*:

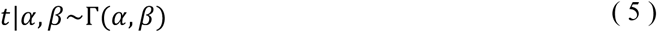

The resulting posterior distribution was another gamma distribution:

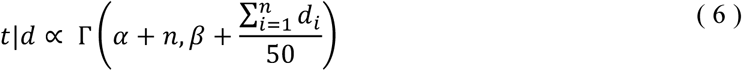

#### Shape (α) and rate (β) of the TMRCA distribution

We modeled the distribution of t as a gamma distribution with shape *α* and rate *β*:

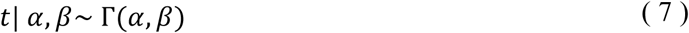

We used the conjugate priors for a gamma distribution rate parameter (*β*) with known shape (*α*), a second gamma distribution:

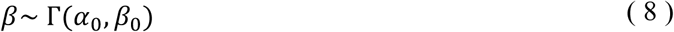

The posterior for *β* then also follows a gamma distribution:

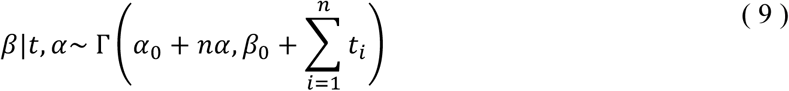

#### Full conditional distributions

To sample from the posterior for each unknown parameter, we derived the full conditional distributions below, ignoring conditionally independent terms. In each iteration of the Gibbs sampler, we sample each parameter from its full conditional distribution, conditioned on the current values of all other parameters. The sampling algorithm is described in the **Supplementary Methods**.

***π***, mixture proportions of *k*:

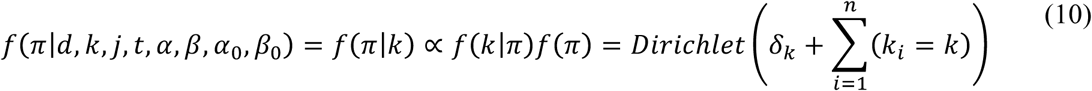

***β***, rate parameter of TMRCA distributions:

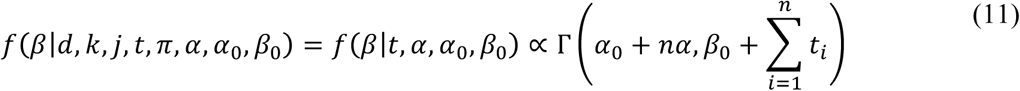

***t***, TMRCA:

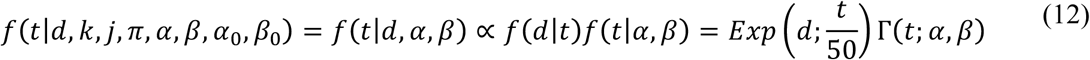

***k***, variant label (IBD or non-IBD partition):

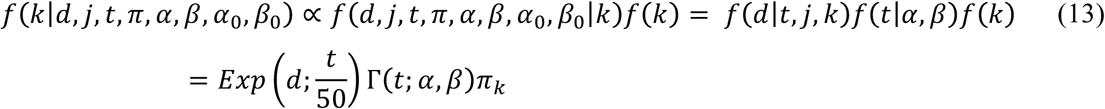

***j***, the partition of non-IBD variants:

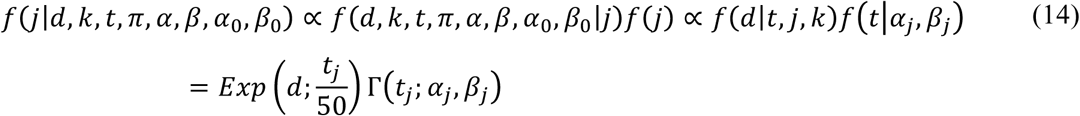

## Results

### Application to simulated genetic data

To evaluate our approach, we applied it to simulated genetic data including non-IBD (recurrent) mutations. Using the forward genetic simulation engine SLiM (Haller and Messer 2017), we generated genomic segments of length 10Mb with uniform mutation and recombination rates (*µ* = 2.5×10^−8^ mutations per site per generation, r = 1×10^−8^ events per site per generation), and no selection, following a European demographic model (Bhaskar et al. 2015). For each simulation, we measured the pairwise obligate recombination distances of recurrent and IBD variants with allele count ≤10 in the 2Mb at the center of each genomic segment. The number of recurrent mutations in these simulations is plotted in **Supplementary Figure 4**.

We applied our Bayesian hierarchical model to the obligate recombination distances from these simulations, and calculated the posterior probability of a variant being non-IBD as the fraction of posterior samples with *k*>1 (**Supplementary Figure 5**). We then evaluated the ability of this posterior estimate to distinguish the IBD and recurrent variants from their obligate recombination distances. The receiver operating characteristic (ROC) curves in **Figure 2** show the relationship between true and false positive rates for allele counts 2-10. The precision and recall of our approach depends on the fraction of variants that are non-IBD (**Figure 2**), with higher recurrent fractions having superior performance.

**Figure 2.**
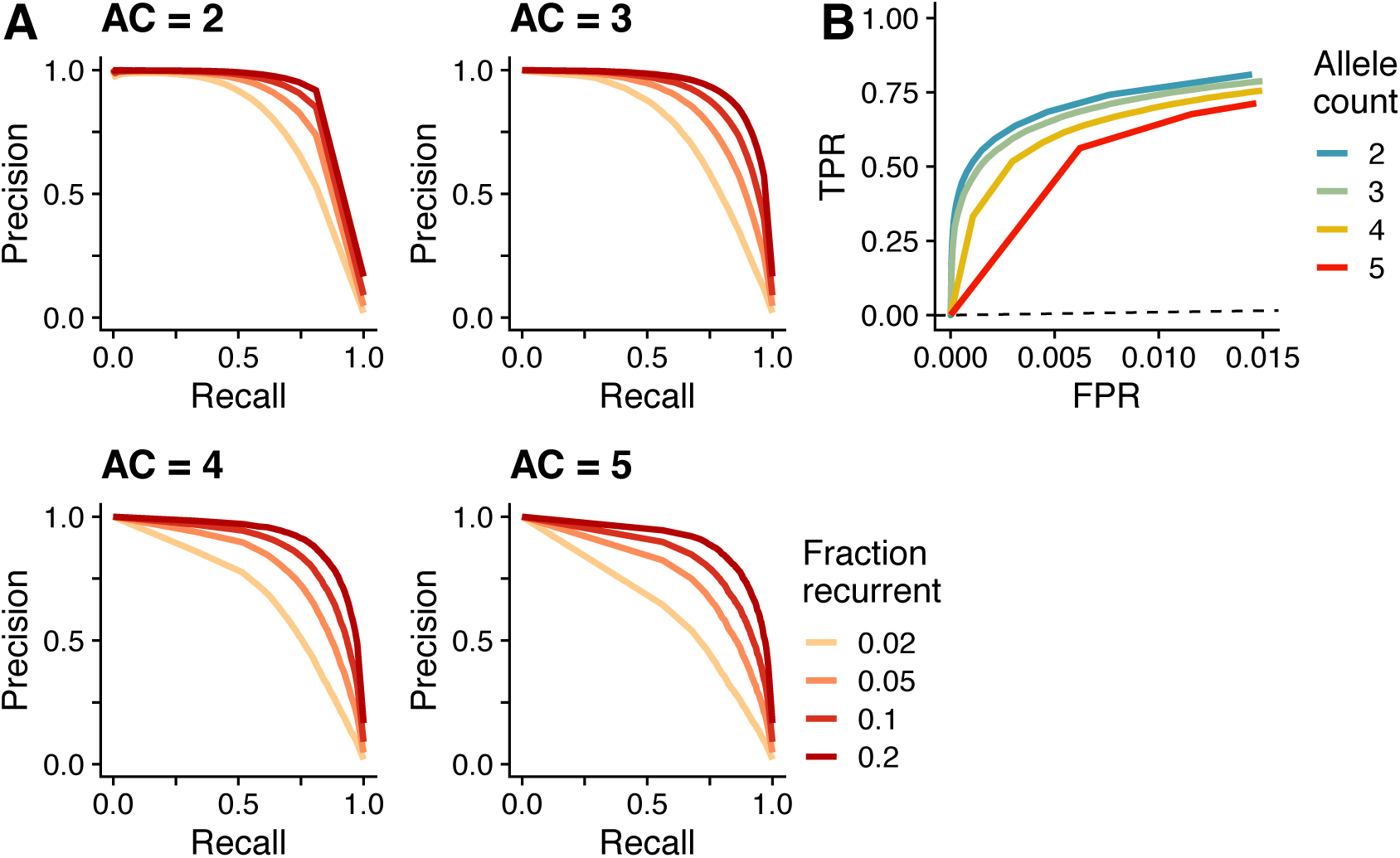
**(A)** Precision-recall plots and **(B)** ROC plots for the Bayesian hierarchical model applied to distinguish recurrent and IBD variants in simulated data. In **A**, each panel represents the application to variants of a given allele count (AC). In **B**, the dashed line represents the identity line.

We next performed a battery of sensitivity studies, simulating population genomics features known to influence patterns of genetic variation that may impact the robustness of our estimates. First, we performed simulations of genomic segments including genes and deleterious mutations, and applied our approach to these simulations to test the effect of selection on our approach to identify non-IBD variants (**Methods**). We found that including background selection had little impact on the power of our approach to identify recurrent mutations (**Supplementary Figure 6; Supplementary Table 1**). We suspect that this may be due to the fact that the rare variants we are interested in are largely quite new (**Supplementary Figure 7**); thus, the difference in recombination distance between IBD and recurrent variants in these simulations is not strongly altered by the presence of weak negative selection.

Next, we evaluated the performance of our approach for simulated variants flanked by 10,000 base pair recombination hotspots, with hotspot recombination rates of 5×10^−6^, 1×10^−6^, or 5×10^−7^ events per base pair per generation (**Methods**). For variants close to a hotspot, we expected a smaller difference between recurrent and IBD allele pairs’ recombination distances, due to a weaker relationship between recombination distance and TMRCA. When we applied our Bayesian hierarchical model to the simulations with hotspots, we found that our power to distinguish IBD and recurrent variants decreased with increased hotspot strength (**Supplementary Figure 8; Supplementary Table 1**).

### Comparison to other approaches to identify non-IBD variants

To provide alternative approaches for comparison and benchmarking purposes, we developed a composite-likelihood approach to identify non-IBD allele pairs, based on coalescent theory of the TMRCA for IBD or recurrent allele pairs in an exponentially growing population (**Supplementary Methods**). While the likelihood-based approach had some power to classify events, this approach performed less well than the Bayesian hierarchical model (**Supplementary Figure 9, Supplementary Table 2**). For allele counts <7, the likelihood-based approach had substantially worse power at the lowest false positive rates, the relevant range for identifying non-IBD variants.

While no genome-wide scalable approach to identify specific non-IBD variants exists to our knowledge, there are recently developed methods that estimate the age of a variant in large scale genome-wide sequencing data (Platt et al. 2019; Albers and McVean 2020). Non-IBD mutations could potentially be identified as outliers in the age estimates of each allele frequency class by these approaches. We estimated simulated variants’ ages using the estimator *runtc* (Platt et al. 2019) (**Methods**). We used these age estimates to distinguish simulated IBD and recurrent variants, and plot the performance of this approach in **Supplementary Figure 10**. We find that the age estimates have limited power to distinguish non-IBD variants, and that *runtc*’s performance at this task – a task we note that it was not explicitly designed for – performs poorly compared to our Bayesian hierarchical approach (**Supplementary Table 2**).

### Application of Bayesian hierarchical model to UK10K sequencing data

We applied our method to identify non-IBD variants in whole-genome sequencing data in 3,621 individuals from the UK10K project (Walter et al. 2015). Individuals from the ALSPAC and TWINSUK studies used here were sequenced to average depth ∼7x and passed the UK10K project quality control filters. We measured the obligate recombination distance for biallelic and multiallelic single nucleotide variants that passed the UK10K quality filters. Based on the decreased performance of our method with increased allele count, we restricted our analysis to variants of allele count less than or equal to 5.

We applied our approach to a mixture of 80% biallelic and 20% multiallelic sites, in order to use multiallelic sites as a positive control for non-IBD mutations. We compared the empirical cumulative distribution of posterior probabilities for multiallelic and biallelic sites, and as expected we observed that multiallelic sites had higher posterior probabilities of being non-IBD (**Supplementary Figure 11**). We used these distributions to determine the threshold of posterior probabilities we called “likely non-IBD” for all biallelic variants at allele counts 2-5, which we then used in downstream analyses.

### Non-IBD variants correlate with local sequence context

To assess the accuracy of our recurrent mutation calls, we took advantage of the relationship between local sequence context and mutation rate (Aggarwala and Voight 2016). Under a Poisson model of mutation, sequence contexts with a higher mutation rate should have a higher probability of recurrent mutation relative to other contexts (i.e., “double hits”). If non-IBD variant calls reflect recurrent mutations, we would expect to see a correlation between the fraction of non-IBD variants and the mutability of sequence contexts. Conversely, if our approach randomly selects a subset of sites rather than true recurrent mutations, we would not expect to see a relationship between sequence context and fraction of sites called recurrent. Using sequence-context estimated polymorphism probabilities calculated from the UK10K dataset, we calculated an expected fraction of recurrent variants for each 5 base-pair (5-mer) sequence context and allele count (**Methods**).

Across all 5-mer sequence contexts, we observed a significant correlation between expected and observed fractions (e.g. Pearson’s correlation = 0.81, P < 10^−100^ for allele count 2; **Figure 3; Supplementary Table 3**). The observed fraction of non-IBD called sites was higher than expected for non-CpG->T contexts, and lower than expected for CpG->T contexts (**Figure 3; Supplementary Table 4**). Within CpG->T contexts, we also observed a significant correlation between expected and observed fractions, though for all contexts the observed fraction of non-IBD calls was less than expected (**Supplementary Figure 12; Supplementary Table 4**). Within non-CpG->T contexts, the correlation between expected and observed was significant for all allele counts except for variants of allele count 5, which have the smallest sample size (**Supplementary Figure 13; Supplementary Table 4**). These results suggest that at sequence contexts with relatively lower polymorphism probabilities, there was a higher rate of non-IBD calls. Non-CpG->T contexts represent 82% of the polymorphic sites tested, but 68% of sites called non-IBD.

**Figure 3.**
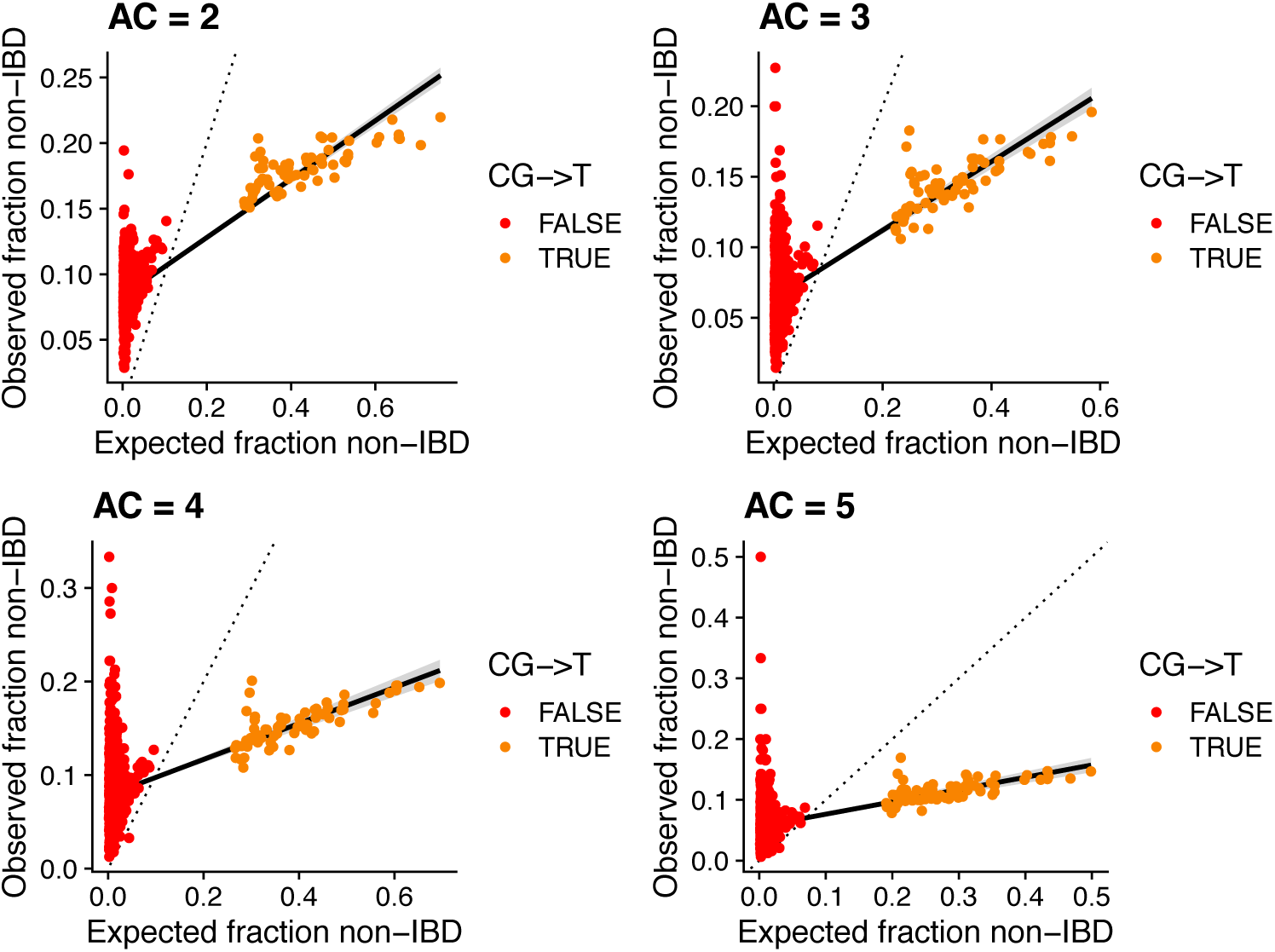
The expected and observed fraction of sites called non-IBD for UK10K variants. Each dot represents a 5-mer sequence context. The expected fraction was calculated from each sequence context’s polymorphism probability. The solid black line is a linear regression line for all sequence contexts, and the dotted line is the identity line.

### Additional genomic annotations correlated with non-IBD variants

Next, to understand which genomic features in addition to local mutation rate are associated with non-IBD variants, we performed a linear regression with the posterior probability of each variant being non-IBD as the response variable (6,763,324 sites; with 665,340 called non-IBD). In separate regressions for each allele count, we included 7-mer polymorphism probabilities, background selection, GC content, replication timing, local recombination rate, distance to a recombination hotspot, germline CpG methylation levels, the variant calling quality measure VQSLOD, and read depth as predictor variables. We transformed the values of each annotation to z-scores, and report the odds ratios and 95% confidence interval for each annotation in **Figure 4 (Supplementary Table 5)**. All annotations were significantly associated with the outcome (P<1×10^−10^). In addition, we performed a logistic regression with IBD/non-IBD calls for each variant as the response variable (**Supplementary Figure 14; Supplementary Table 5**). We also performed regressions with CpG->T sites only (**Supplementary Figure 15, Supplementary Table 5**). Below, we highlight the annotations included as predictors, our prior hypotheses about their relationships with recurrent mutations, and the results of the regression models.

**Figure 4.**
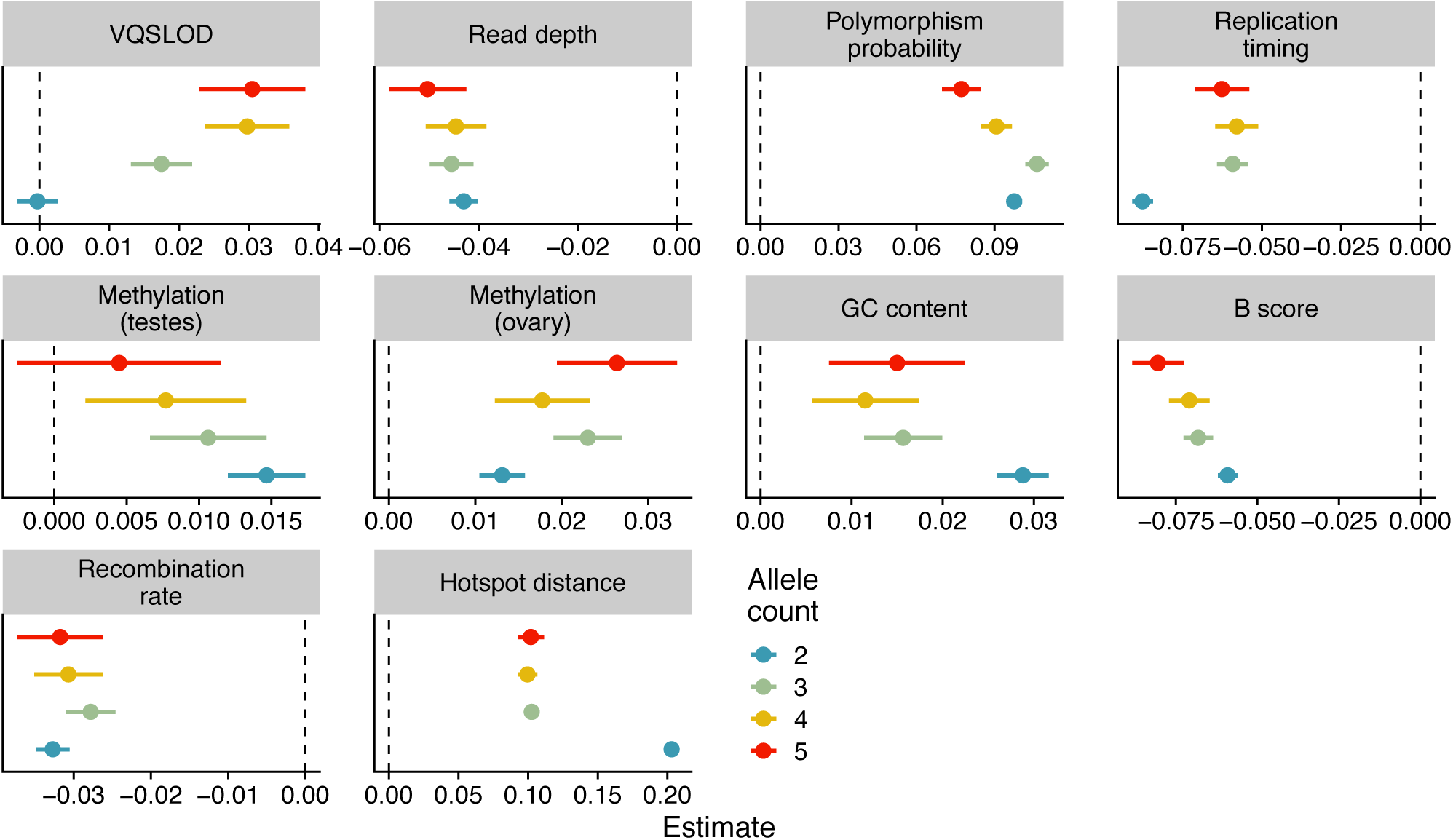
Linear regression of genomic annotations (predictor variables) vs. posterior probability of being non-IBD (outcome) for all variant sites, grouped by allele count. Dot colors represent allele count, and a separate regression was run for variants of each allele count. Each dot’s position denotes its beta coefficient estimate, with error bars representing beta ± 1.96*standard error. The vertical dashed line represents a beta estimate of zero. Hotspot distance: physical distance to nearest recombination hotspot z-score; Recombination rate: local recombination rate z-score; B score: McVicker’s B statistic z-score; Replication timing: replication timing z-score; GC content: local GC content z-score; Methylation (ovary): ovary CpG methylation z-score; Methylation (testes): testes CpG methylation z-score; Read depth: read depth z-score; VQSLOD: variant quality z-score.

#### Polymorphism probability

As shown in our analysis of expected vs. observed recurrent fraction for 5-mer sequence contexts above, polymorphism probability was strongly positively correlated with non-IBD status. As previous work has shown that a 7-mer model explains additional variation in genetic variation over a 5-mer model (Aggarwala and Voight 2016), we find that a 7-mer polymorphism probability calculated in UK10K in the logistic regression model was associated with our non-IBD calls.

#### GC content

GC content varies across the human genome, and is correlated with gene content, repetitive elements, DNA methylation, recombination rates, and substitution probabilities (Arndt et al. 2005). In our regression model, increased local GC content (measured at a 1kb scale) was associated with increased probability of a variant being called non-IBD.

#### Replication timing

Later replication timing has been linked to higher rates of de novo mutations in the human genome, specifically in the offspring of relatively younger fathers (Francioli et al. 2015). Our regression model with replication timing estimates (Koren et al. 2012) was consistent with these results, with variants in late replicating regions significantly more likely to be called as recurrent (positive replication timing values mean earlier replication).

#### Background selection

We included B-values (McVicker et al. 2009), a measure of background selection, or purifying selection due to linkage with deleterious alleles. Lower B-values indicate a lower fraction of neutral variation in a region, i.e., stronger background selection. We expected that increased background selection would be associated with increased recurrent mutation, as linkage to deleterious alleles would result in variants being removed from the population and thus present at lower frequencies. Recurrent mutations would then be more likely to be present as they effectively shift the site frequency spectrum towards more rare alleles. Our results are consistent with this expectation, with an odds ratio less than one for B-values.

#### Local recombination rate and distance to recombination hotspots

The results of our simulations suggested that we have lower power to identify recurrent variants located near a recombination hotspot (**Supplementary Figures 8, 9**). Indeed, we observed that both an increased local recombination rate and a shorter distance to a recombination hotspot were correlated with a lower probability of a site being called as recurrent.

#### Methylation levels at CpG sites

Spontaneous deamination of 5-methylcytosine at CpG sites results in a substantial increase in C-to-T transition mutations. We included CpG methylation levels measured in testes and ovaries in our model, expecting that CpG sites with higher methylation levels are more likely to spontaneously deaminate, increasing mutation rates generally and thus increase recurrent mutation probabilities. Methylation levels in testes and ovaries were correlated (Pearson’s correlation coefficient = 0.27, P<2×10^−16^), but we noted that increased methylation in both tissue types independently predicted an increased posterior probability of a variant being non-IBD.

#### VQSLOD and read depth

We observed a significant relationship between sequencing quality, measured both by read depth and variant quality score, and the probability of a site being non-IBD. Under a simple model for genotyping error, where errors are distributed randomly (without respect to haplotype), this result suggests that our approach also identifies some number of genotyping errors in regions of low read depth or sequencing quality.

### Non-IBD calls and gene conversion events

As non-crossover gene conversions are thought to be more frequent than de novo mutations in the human genome (Halldorsson et al. 2016), we expect that a subset of our non-IBD variant calls reflect gene conversion events. After a non-crossover gene conversion event encompassing a rare variant, the copied allele resides on the existing haplotype background of the acceptor chromosome, which may reduce the surrounding shared IBD segment. We devised a heuristic to identify likely gene conversions, based on the intuition that two non-IBD variants in close physical proximity in the same individuals are more likely to reflect variants copied along a gene conversion tract, rather than two independent recurrent point mutations. If a gene conversion tract contains only a single rare variant, this signature would be indistinguishable from a recurrent point mutation with our approach. Furthermore, if a gene conversion contained no rare variants, it would not be identified in our analysis as a potential recurrent mutation or gene conversion.

Limiting our results to tracts less than 1kb with 2 or more non-IBD variants present in the same individuals, we identified 42,203 variants within 18,971 putative gene conversion tracts, representing 6.3% of non-IBD variants (**Supplementary Figure 16**). We performed logistic regression with all non-IBD variants labeled as potential gene conversions or not as the outcome, and the genomic annotations listed above as predictor variables (**Figure 5, Supplementary Table 6**). We additionally included the posterior probability of a variant being non-IBD as a predictor variable. Compared to non-IBD variants not in putative gene conversion tracts, these variants were associated with lower polymorphism probability, higher variant quality score, increased posterior probability of being non-IBD, smaller distance to a recombination hotspot, and lower recombination rate. We also observed a GC bias in putative gene conversion variants, as measured by the fraction of variants containing an A->C/T->G or A->G/T->C mutation (37% in putative gene conversions vs. 27% in all other non-IBD variants; P < 10^−100^; Fisher’s exact test).

**Figure 5.**
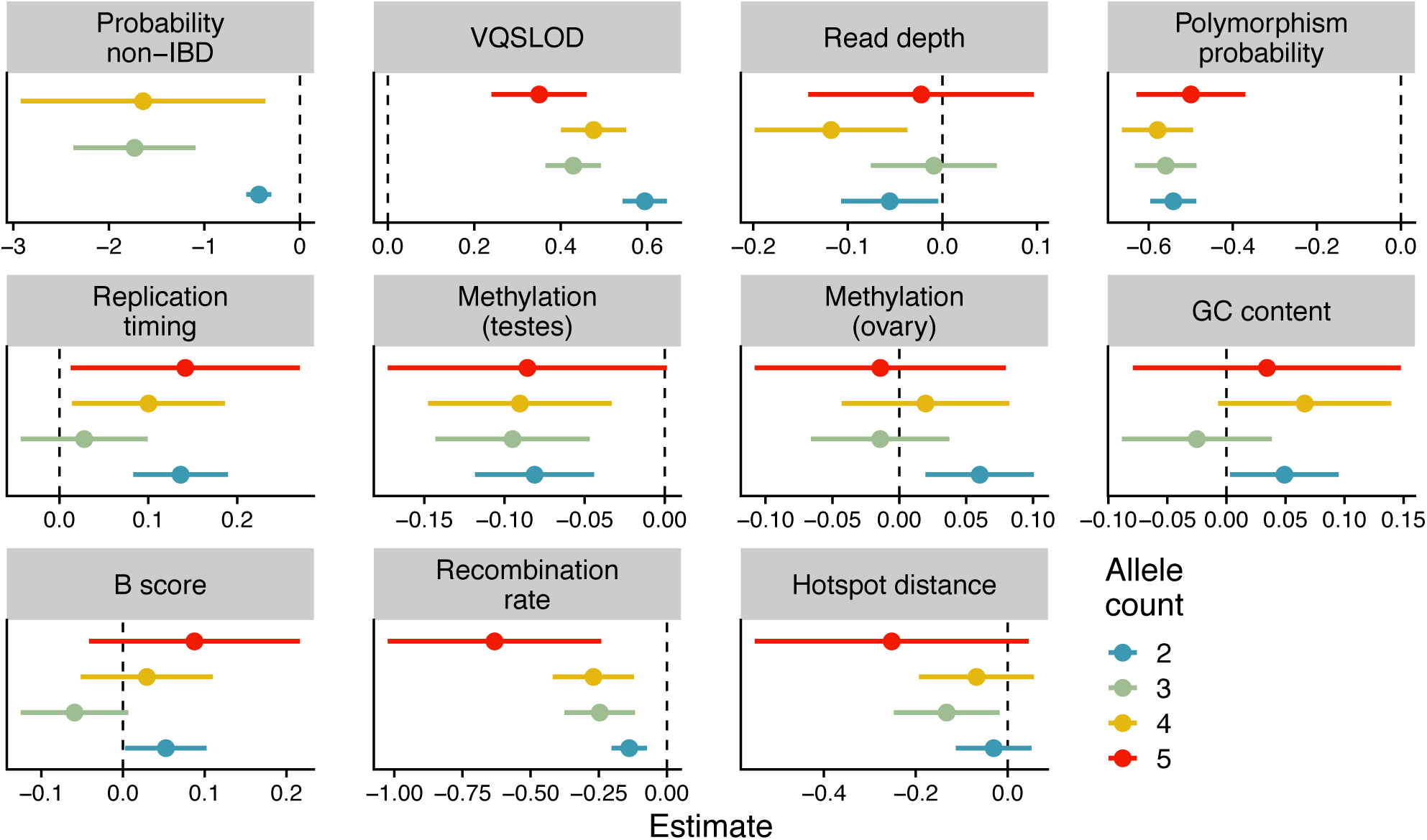
Results of a logistic regression using genomic annotations to distinguish putative gene conversions from other non-IBD variants. Separate regressions were performed for variants of each allele count. The annotation of variants’ probability of being non-IBD for allele count 5 was left off to improve the visualization (Estimate: -7.0; 95% CI: -16.4 - 2.4). Dot colors represent allele count. Each dot’s position denotes its beta coefficient estimate, with error bars representing the 95% confidence interval (beta ± 1.96*standard error). The vertical dashed line represents a beta estimate of zero. Hotspot distance: physical distance to nearest recombination hotspot z-score; Recombination rate: local recombination rate z-score; B score: McVicker’s B statistic z-score; Replication timing: replication timing z-score; GC content: local GC content z-score; Methylation (ovary): ovary CpG methylation z-score; Methylation (testes): testes CpG methylation z-score; Read depth: read depth z-score; VQSLOD: variant quality z-score; Probability non-IBD: posterior probability of variant being non-IBD.

### Rescaling the site frequency spectrum with recurrent mutations

With our set of high-confidence non-IBD variants, we rescaled the site frequency spectrum for very rare variants. Taking into account the power of our approach on simulated data, we plot the original and rescaled SFS for variants with allele count <5 in **Figure 6A** (**Methods**). Rescaling the site frequency spectrum resulted in a 3% increase in the fraction of singleton variants, from 46.6% to 49.6%. As expected, the majority of this shift is due to the relatively large fraction of CpG->T variants that were called as non-IBD (**Figure 6B**). For CpG->T variants alone, the fraction of singleton variants increased from 43.6% to 49.9%. We note that this rescaling is incomplete, as we identified non-IBD variants at only allele counts 2 to 5 (representing 38% of non-singleton variants in UK10K).

**Figure 6.**
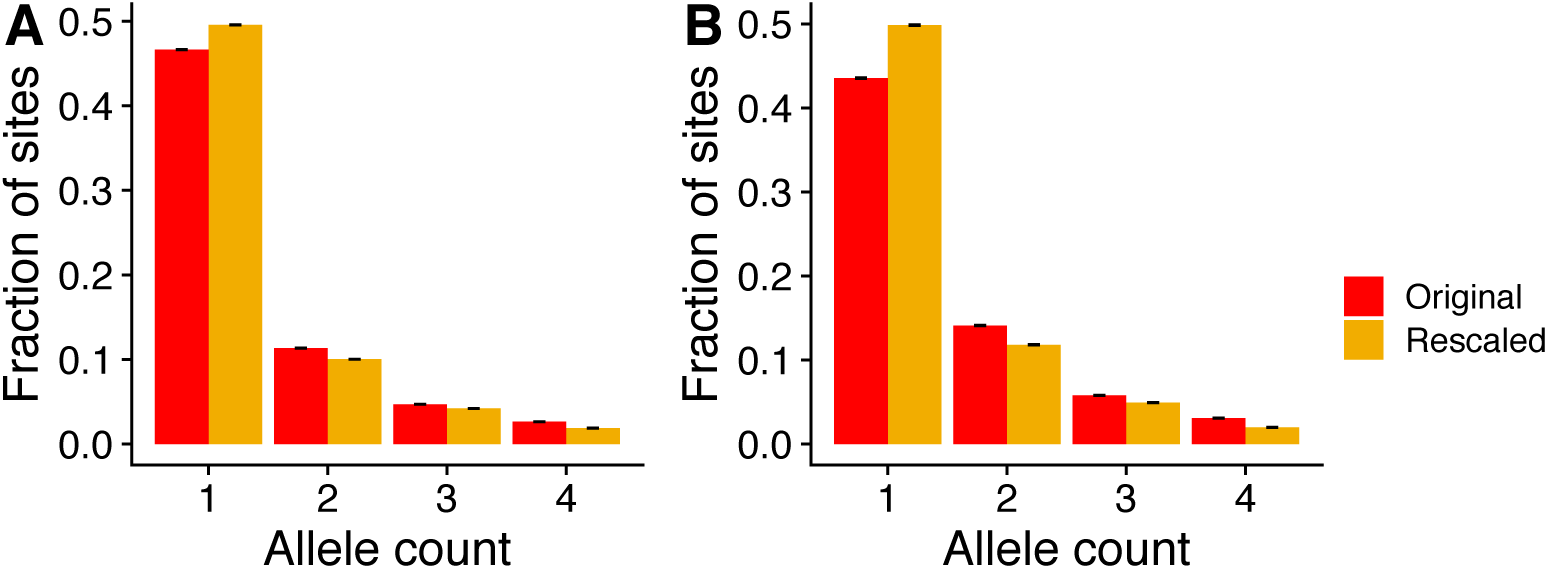
The site frequency spectrum for variants of allele count <5, before and after rescaling to incorporate non-IBD variants. **(A)** The original and rescaled SFS for all variants. **(B)** The original and rescaled SFS for CpG->T variants only.

## Discussion

We describe a novel approach designed to specifically identify non-IBD variants in whole genome sequencing data by leveraging the difference in the obligate recombination distance between rare IBD and non-IBD variants. Our approach uses a Bayesian hierarchical model and Gibbs sampling to jointly infer the TMRCA distributions of these two scenarios and identify variants with a high posterior probability of being non-IBD. In simulated data, we find that the posterior probabilities of a variant being non-IBD can discriminate between IBD and recurrent mutations for variants up to allele count 5 in a population sample of 3,621 individuals.

Our approach assumes that we do not have phase information for individuals, i.e. we do not assign each variant in an individual to a maternally or paternally inherited chromosome. If we had accurate phase information for rare variants, such as from long-read sequencing data, or ‘hard-phase’ calls from paired-end sequencing libraries, we could more accurately measure recombination breakpoints. This could potentially improve the accuracy of our method by eliminating the measurement error caused by using the obligate recombination distance.

We focused on identifying non-IBD variants for allele counts of 5 or less, as the performance of our method decreases with increasing allele count. Additionally, the computational burden of sampling from the marginal posterior distributions increases exponentially with increasing allele count. With a larger sample size, the frequency of a variant at a given allele count will decrease, while the computational complexity remains the same. Thus, we expect that applying our method to even larger sequencing datasets will improve its performance.

The negative correlation we observed between local recombination rate and the probability of a site being called non-IBD suggests that our method is confounded by local recombination rate. In simulated data, we also observed that we had lower power to identify recurrent mutations in close proximity to recombination hotspots. We also note that we found a significant relationship between sequencing quality, measured by read depth and variant quality score, and the probability of a site being called non-IBD. The signature of a non-IBD variant used here could also be that of a genotyping error, as genotyping errors also may occur on any random haplotype background in a population. This could be a potential application of our method, as a way to identify genotyping errors in large scale sequencing datasets. Our current recommendation to overcome this issue is to remove variants of low quality until this relationship is not significant. However, distinguishing genotyping errors from true non-IBD variants remains an important problem.

## Materials and Methods

### Forward genetic simulations with SLiM

We used the software program SLiM version 2.5 (Haller and Messer 2017) for forward genetic simulations. We used the following European demographic model (Bhaskar et al. 2015): an ancestral population size of 10,000 with a burn-in period of 100,000 generations; a population bottleneck to 200 individuals at generation 200; population size rebounds to 10,000; a second bottleneck to 500 individuals at generation 4,280; population size rebounds to 5,800; exponential growth starting at generation 4,870 at 3.89% per generation; random sampling of 3,621 individuals at generation 5,000. SLiM simulations had a uniform mutation rate of 2.5×10^−8^ mutations per base pair per generation. We identified recurrent mutations as base positions with two or more unique mutations. We performed 1,000 simulations with uniform recombination rate of 1×10^−8^ events per base pair per generation, and additional 100 simulations each with recombination hotspots of r = 5×10^−6^, 1×10^−6^, or 5×10^−7^. Each 10Mb simulated genomic segment had two hospots of of length 10,000 bp flanking the central 2Mb of the segment.

For forward genetic simulations with selection, we generated 10Mb genomic segments using a recipe from the SLiM manual (Haller and Messer 2017) with the following procedure: 1) sample non-coding region; 2) sample exon; 3) sample intron and exon pairs in a loop with 20% probability of stopping after each pair; 4) repeat steps 1-3 while chromosome length < 10Mb; 5) sample final non-coding region. Exonic mutations were synonymous or non-synononymous at a ratio of 1:2.31, and 10% of non-synonymous mutations were neutral. Deleterious non-synonymous mutations’ selection coefficients were sampled from a gamma distribution with mean -0.03 and shape 0.207. Exon lengths were sampled from a lognormal distribution with mean log(50) and standard deviation log(2). Non-coding regions were neutral and their lengths were sampled from a uniform distribution between 100 and 5000. Intronic mutations were neutral and intron lengths were sampled from a lognormal distribution with mean log(100) and standard deviation log(1.5).

### UK10K dataset

We applied our method to identify recurrent mutations in whole-genome sequencing data in 3,621 individuals from the UK10K project (Walter et al. 2015). These individuals were sequenced to average depth 7x, passed the UK10K project quality control filters, and come from the ALSPAC and TWINSUK studies. We measured the recombination distance for biallelic and multiallelic single nucleotide variants that passed the UK10K quality filters that were present at allele count ≤ 10 in these individuals.

### Measuring the obligate recombination distance

For simulated data, we generated diploid genotypes by randomly combining pairs of haploid genomes, and calculated the recombination distances for variants within the central 2Mb of each 10Mb genomic segment. In both simulated and UK10K data, we measured the obligate recombination distances for variants with allele count ≤ 10. For each pair of carriers, we identified the nearest variant upstream and downstream with opposite homozygote genotype, i.e. where one individual has genotype 0 and the other has genotype 2 (**Supplementary Figure 3**). We then converted the physical distance to a genetic distance using a genetic map. For UK10K, we used a 1000 Genomes Project CEU genetic map (The 1000 Genomes Project Consortium 2012), and for simulated data we used a uniform map with *r* = 1×10^−8^ events per site per generation, or a variable map for simulations including recombination hotspots.

### Applying the Bayesian hierarchical model

To apply our model to simulated or UK10K recombination distances, we first generated an estimate of the beta parameter for non-IBD variants from multiallelic sites. Using Gibbs sampling on non-IBD allele pairs from multiallelic variants, we used a simplified version of the hierarchical model where we sampled the TMRCA for each allele pair and the beta parameter in each Gibbs iteration. We repeated this procedure to estimate beta for a range of alpha values from multiallelic sites’ recombination distances. To test if the choice of alpha affected our ability to discriminate IBD and non-IBD variants, we applied the model with different non-IBD alpha/beta values to UK10K variants on chromosome 22. The posterior estimates of k were highly correlated across values of alpha (alpha=20 vs. alpha=40, **Supplementary Table 7**). When applying the full model to data, we used alpha = 10 for IBD allele pairs, with alpha = 40 and the corresponding value of beta inferred from multi-allelic sites (beta = 0.0859) for non-IBD allele pairs. We ran 10,000 iterations of the Gibbs sampler for each run of the model, thinned the chains until autocorrelation was below 0.01, and assessed convergence of the chains by comparing the thinned samples from the first and second half of the chain via a Wilcoxon rank-sum test. A chain was determined to have converged if the Wilcoxon test P-value was > 0.05.

We parallelized the application of our model by breaking down the genome into 10Mb segments, rather than including all variants of a given allele count in a single run of the Gibbs sampler. To test the effect of the number of variants included in a Gibbs sampling run, we applied the model to 10Mb segments on chromosome 22 and to all variants on chromosome 22 together. For smaller allele counts with thousands of variants in each segment, we observed no effect, but for larger allele counts we did see an effect of applying the model to small numbers of variants. Thus, for allele counts >5, we grouped segments together until at least 1000 variants were included in each run of the model.

### Variant age estimation with runtc

The *runtc* software was downloaded from https://github.com/jaredgk/runtc (Platt et al. 2019). The output from 100 simulations from SLiM with uniform recombination and mutation rates was converted to VCF format, and then runtc was applied to the vcf files with the commands --k-range 2 10 --rec 1e-8 --mut 2.5e-8.

### Area under the ROC curve (AUC)

For all ROC curves from simulated data, we calculated the area under the curve as:

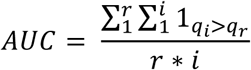

where *r* and *i* represent recurrent and IBD variants, and *q*_*r*_ and *q*_*i*_ the values of the statistic being evaluated. For each AUC, we calculated a confidence interval by generating 10,000 bootstrap samples of 5,000 variants (with the same ratio of IBD:recurrent variants as the simulated sample). We then sorted the 10,000 AUC estimates and took the 2.5th and 97.5th percentiles to get a 95% confidence interval.

### Calculating an expected fraction of recurrent mutations from polymorphism probabilities

As a proxy for the mutation rate, we estimated the polymorphism probability for 5-mer sequence contexts (i.e., the focal base and two bases up and downstream) as the fraction of sites with that context that were variable in the UK10K dataset. These polymorphism probabilities were highly correlated with those calculated previously with the 1000 Genomes dataset (Aggarwala and Voight 2016) (Pearson’s correlation = 0.99, P<10^−100^), with a higher fraction of polymorphic sites in the UK10K for each context due to the larger sample size (**Supplementary Figure 5**).

To predict the fraction of sites that should be called recurrent based on sequence context polymorphism probabilities, we used a simple Poisson model of mutation. With the polymorphism probability for a context as the Poisson rate parameter *λ* and the number of mutations at a site *H*, the probability of a recurrent mutation is the probability of two or more mutations at a site:

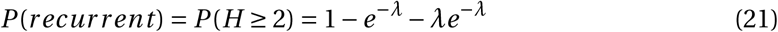

As we are only considering sites where there has been at least one mutation event, i.e. polymorphic sites, the probability of a recurrent mutation at a site is then:

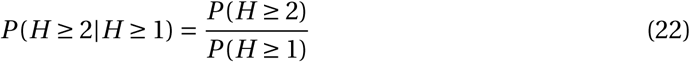

We calculated this probability for each 5-mer sequence context. We then calculated the expected fraction by scaling the overall fraction of sites called non-IBD by each context’s probability of a recurrent mutation, relative to all the other contexts.

We used 5-mer sequence contexts for this analysis so that we would have a reasonable number of variants classified as IBD or not for each sequence context. If we had used 7-mer sequence contexts, some contexts would have too few variants to calculate the proportion called non-IBD. For the regression models to predict non-IBD variants using multiple genomic annotations, we used 7-mer sequence contexts, as there is significant mutation rate variation even within 5-mer contexts (Aggarwala and Voight 2016).

### Identifying putative gene conversions

Within the set of variants called as non-IBD, we called putative gene conversion tracts that contained 2 or more variants that were: 1) present in the same individuals, 2) at the same allele count, 3) within 1kb of each other.

### Genomic annotation datasets

We used the B statistic (McVicker et al. 2009) (downloaded from http://www.phrap.org/othersoftware.html) to measure background selection, which estimates the proportion of neutral variation in a region. VQSLOD and read depth were extracted from the UK10K VCF files. We used a recombination rate map estimated for Europeans from the 1000 Genomes Project, downloaded from http://ftp.1000genomes.ebi.ac.uk/vol1/ftp/technical/working/20130507_omni_recombination_rates (The 1000 Genomes Project Consortium 2012). We used human recombination hotspots identified in the HapMap project (The International Hapmap Consortium 2007), and downloaded from https://github.com/auton1/Campbell_et_al. Replication timing data was obtained from (Koren et al. 2012). CpG methylation levels were downloaded from https://www.ncbi.nlm.nih.gov/geo/ using accession numbers GSM1010980 (ovary), and GSM1127119 (testis).

### Rescaling the SFS with non-IBD mutations

Starting with the SFS calculated from all UK10K biallelic sites included in our study, for allele counts 2-5 for CpG->T and all other mutation types we calculated the fraction called as non-IBD. We then divided this fraction by the power of our method, estimated by the percent of multiallelic sites identified as non-IBD at the chosen posterior threshold. From this fraction of non-IBD sites for the two mutation types, we apportioned the non-IBD mutations into lower allele counts based on the relative frequency of allele counts 1-5. For example, to determine what fraction of non-IBD 4-ton variants would be assigned partition 1:3 vs. 2:2, we used the relative frequencies:

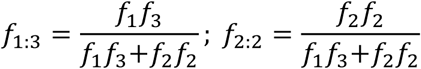

Where *f*_1:3_ is the relative frequency of the 1:3 partition, and *f*_1_ is the frequency of singletons in the original SFS. In the rescaled SFS, the number of singletons increased by the number of variants of allele count 2-5 that were identified with partition 1:(n-1); the number of doubletons decreased by the number of doubletons that were identified as recurrent, and increased by the number of variants of allele count 3-5 that had partition 2:(n-2); and so on through allele count 4. Allele count 5 was excluded from the rescaled SFS plots because we did not identify recurrent variants at allele counts greater than 5.

## Acknowledgements

This work was supported by the US National Institutes of Health (grant numbers DK101478 to B.F.V. and T32 GM008216 for K.E.J.) and a Linda Pechenik Montague Investigator award (to B.F.V.). This study makes use of data generated by the UK10K Consortium, derived from samples from ALSPAC and TWINSUK. A full list of the investigators who contributed to the generation of the data is available from www.UK10K.org. Funding for UK10K was provided by the Wellcome Trust under award WT091310.

## Statement of Work

K.E.J. and B.F.V. conceived of the experiments, designed the methodology, analyzed the data, and wrote the manuscript. B.F.V. supervised the work.

## Statement of Competing Interest

The authors declare no competing interest.

## Data availability

The Gibbs sampler for the Bayesian hierarchical model is available as an R package at github.com/kelsj/ibdibsR. The hierarchical model input data (pairwise obligate recombination distances) and output (posterior probabilities) from simulations and UK10K are available at http://coruscant.itmat.upenn.edu/data/Johnson_Voight_Sims_UK10K_PP_nonIBD.tar.gz.

## Notes

http://coruscant.itmat.upenn.edu/data/Johnson_Voight_Sims_UK10K_PP_nonIBD.tar.gz

### Competing Interest Statement

The authors have declared no competing interest.

