## Supplementary Methods & Figures for "Identifying non-identical-by-descent rare variants in population-scale whole genome sequencing data"

K.E. Johnson & B.F. Voight

**Supplementary Methods** ... **2**
*Theoretical distributions of TMRCA for recurrent or IBD alleles* ... 2
*Gibbs sampling algorithm for the Bayesian hierarchical model* ... 3
*Likelihood-based model to identify non-IBD variants* ... 3
*Application of likelihood-based model simulated data* ... 6

**Supplementary Table Descriptions** ... **7**

**Supplementary Figures** ... **9**
Supplementary Figure 1. Theoretical distributions of the time to most recent common ancestor. ... 9
Supplementary Figure 2. Theoretical distributions of the recombination distance. ... 9
Supplementary Figure 3. Measuring the obligate recombination distance with unphased diploid genotypes. ... 10
Supplementary Figure 4. Counts of IBD and recurrent variants in 1000 simulations. ... 10
Supplementary Figure 5. Posterior probabilities of simulated IBD or recurrent variants being non-IBD. ... 11
Supplementary Figure 6. ROC curves of the Bayesian hierarchical model's ability to distinguish IBD and non-IBD variants from simulations with or without selection. ... 11
Supplementary Figure 7. The distribution of allele ages for IBD or recurrent mutations in simulations with and without selection. ... 12
Supplementary Figure 8. ROC curves of the Bayesian hierarchical model's ability to distinguish IBD and non-IBD variants from simulations with recombination hotspots. ... 13
Supplementary Figure 9. ROC curves comparing the performance of the Bayesian hierarchical model to a theoretical likelihood-based model. ... 13
Supplementary Figure 10. ROC curves comparing the performance of the Bayesian hierarchical model to age estimates from *runtc*. ... 14
Supplementary Figure 11. The distribution of posterior probabilities of being non-IBD for biallelic vs. multiallelic variants from UK10K. ... 14
Supplementary Figure 12. The expected and observed fraction of sites called non-IBD for UK10K variants at CpG sites. ... 15
Supplementary Figure 13. The expected and observed fraction of sites called non-IBD for UK10K variants at non-CpG sites. ... 16
Supplementary Figure 14. Results of a logistic regression using genomic annotations to predict non-IBD variant calls. ... 17
Supplementary Figure 15. Results of linear or logistic regression using genomic annotations to predict non-IBD variant calls for CpG variants. ... 18
Supplementary Figure 16. Distribution of putative gene conversion tract lengths. ... 19

### Supplementary Methods

#### *Theoretical distributions of TMRCA for recurrent or IBD alleles*

For two alleles derived from recurrent mutations, the probability distribution of the TMRCA is that of any two random individuals selected from a population. For a neutrally evolving population of constant size $N_e$, this is the probability that the two individuals do not coalesce for *t-1* generations, and then coalesce at generation *t*:

$$f(t) = \left(1 - \frac{1}{2N_e}\right)^{t-1} \frac{1}{2N_e} \quad (1)$$

For a population undergoing exponential growth, or with any other non-constant population size function *N(t)*, we simply replace the constant population size value $N_e$ with *N(t)*:

$$f(t) = \frac{1}{2N(t)} \prod_{i=1}^{t-1} \left(1 - \frac{1}{2N(i)}\right) \quad (2)$$

Assuming exponential growth, and using the continuous time approximation to the coalescent, the probability of a coalescent event between *t* and *t+dt* is (Slatkin and Hudson 1991):

$$f(t)dt = \frac{e^{rt}}{N_0} exp\left(-\frac{e^{rt} - 1}{N_0 r}\right) dt \quad (3)$$

For two IBD alleles with a given allele frequency, to calculate *f(t)* we must account for the fact that the IBD alleles coalesce before all other chromosomes in the sample. Slatkin gave the following approximation for an allele's age, when it is at frequency *x*, in a population of changing size (Slatkin 2000):

$$f(t) \approx C\bar{A}(t)(\bar{A}(t) - 1)(1 - x)^{\bar{A}(t)-2} \quad (4)$$

Where *C* is a normalizing constant, and $\bar{A}(t)$ is the approximate expected value of the number of ancestral lineages at time *t* in the past with sample size *n*:

$$\bar{A}(t) \approx \frac{n}{1 + \frac{n\tau(t)}{2}} \quad (5)$$

*τ(t)* is the population size as a function of time *t,* for an exponentially growing population with growth rate *r* and population size $N_0$ at time *t* = 0.

$$\tau(t) = \int_0^t \frac{dt}{2N(t)} = \frac{e^{rt} - 1}{2N_0 r} \quad (6)$$

Assuming a star-shaped genealogy, we can use this approximate age of a (rare) mutation as the TMRCA for two IBD alleles.

#### *Gibbs sampling algorithm for the Bayesian hierarchical model*

We begin by randomly assigning starting values of each parameter. Then, in each iteration, we sample from the full conditionals in the following steps:

1. Sample $\pi$ from $\pi|k$
2. Sample $\beta$ from $\beta|t,\alpha,\alpha_0,\beta_0$
3. For each potential value of $k$:
   a. Sample $t_k$ from $t|d,k,j,\alpha,\beta$
   b. Sample $j_k$ from $j|d,k,t,\alpha,\beta$
   c. Calculate $P(k=k|d,t_k,j_k,\alpha,\beta)$
4. Sample $k$ from $P(k=k|\ldots)$; accept corresponding $t_k$, $j_k$
5. Go back to step 1.

#### *Likelihood-based model to identify non-IBD variants*

We can calculate the probability of the observed recombination distance $d$ by integrating over all possible values of the TMRCA:

$$f(d,t) = f(d|t)f(t) \tag{7}$$

$$f(d) = \int_0^\infty f(d|t)f(t)dt \tag{8}$$

Following an exponential distribution with rate parameter $\lambda=t/(50\text{ cM})$ for the distance to the nearest recombination event, we have the following expression for the probability of distance $d$ given TMRCA $t$:

$$f(d|t) = \lambda e^{-\lambda d} \tag{9}$$

Thus, for any pair of IBD alleles, we can calculate the probability of the observed recombination distance:

$$f(d_{IBD}) = \int_0^\infty f(d|t)f(t)dt = \int_0^\infty \lambda e^{-\lambda d}\bar{A}(t)(\bar{A}(t)-1)(1-x)^{\bar{A}(t)-2}dt \tag{10}$$

With $\lambda=t/(50\text{ cM})$ and $\bar{A}(t)$ as defined in Equation 5 above.

For recurrent variants with allele count greater than 2, pairs of alleles could be IBD or not depending on the partition of the alleles. For example, a recurrent variant with an allele count of 4 could be a pair of IBD doubletons (2:2 partition), or a singleton and an IBD tripleton (1:3 partition). For IBD allele pairs within a recurrent variant, their distance probability distribution follows Equation 10 above, but with an allele count based on the partition of the recurrent mutation. For non-IBD allele pairs, we instead use the probability density for *f(t)* for a random pair of alleles from an exponentially growing population (Slatkin and Hudson 1991):

$$f(d_{rec}) = \int_0^\infty f(d|t)f(t)dt = \int_0^\infty \lambda e^{-\lambda d} \frac{e^{rt}}{N_0} exp\left(-\frac{e^{rt}-1}{N_0 r}\right) dt \tag{11}$$

For a variant in data, we can calculate the likelihood of each allele pair being IBD or recurrent from the above probability distributions, and the composite likelihood as the product of the likelihood for each allele pair (assuming independence between allele pairs). E.g., to calculate the likelihood of an IBD mutation with *k* allele pairs, we take the product of the likelihood of all the recombination distances:

$$P(IBD) = \prod_k f(d_{IBD}) \tag{12}$$

For recurrent mutations, we must take the set of potential partitions *P* into account. We do this by weighting the probability for each possible partition by the relative frequencies of its components in the observed site frequency spectrum. For example, a recurrent 4-ton could be comprised of a singleton and a tripleton (a 1:3 partition) or two doubletons (a 2:2 partition). We calculate the weights $w^p$ for these partitions as:

$$q_{tot} = q_1 q_3 + q_2^2 \tag{13}$$

$$w^{1:3} = \frac{q_1 q_3}{q_{tot}} \tag{14}$$

$$w^{2:2} = \frac{q_2^2}{q_{tot}} \tag{15}$$

Where $q_c$ is the observed frequency of an allele with count *c*. We could then calculate the likelihood of an observed recombination distance for a recurrent allele pair as the weighted sum of the likelihoods from each partition *p*:

$$P(rec) = \sum_P w^p f(d_{rec}^p) \; for \; p \in P \tag{16}$$

Instead of calculating the exact probability of each allele pair's recombination distances, we found that counting the number of allele pairs *m* with distances longer than some threshold *l* was a more powerful metric to distinguish IBD and recurrent variants. This approach also requires less computation, as we need only calculate the following probability of a recombination distance being greater than the threshold under recurrent or IBD scenarios once.

$$P(D \geq l) = \int_{l}^{\infty} f(D)dD = \int_{D=l}^{\infty}\int_{t=0}^{\infty} f(D|t)f(t)dtdD \tag{17}$$

For IBD mutations, all allele pairs are IBD, so we can calculate the likelihood of the observed distance using the TMRCA distribution for IBD allele pairs:

$$P_{IBD} = P(D \geq l|IBD) = \int_{D=l}^{\infty}\int_{t=0}^{\infty} f(D|t)f(t_{IBD})dtdD \tag{18}$$

For recurrent mutations, we calculate the likelihood of the observed distance with the recurrent or IBD allele pair TMRCA distributions, and the overall likelihood is a weighted sum based on the fraction of recurrent ($w^{p}{}_{rec}$) and IBD ($w^{p}{}_{IBD}$) allele pairs for each partition:

$$\begin{aligned} P_{rec} &= P(D \geq l|rec) \\ &= \sum_{P}\left\{w_{IBD}^{p}\int_{D=l}^{\infty}\int_{t=0}^{\infty} f(D|t)f(t_{IBD})dtdD \right. \\ &\quad \left. + w_{rec}^{p}\int_{D=l}^{\infty}\int_{t=0}^{\infty} f(D|t)f(t_{rec})dtdD\right\} for\ p \in P \end{aligned} \tag{19}$$

Thus, we integrate over the following expressions for IBD or recurrent allele pairs:

$$f(D|t)f(t_{IBD}) = \lambda e^{-\lambda D}\bar{A}(t)(\bar{A}(t)-1)(1-x)^{\bar{A}(t)-2} \tag{20}$$

$$f(D|t)f(t_{rec}) = \lambda e^{-\lambda D}\frac{e^{rt}}{N_0}exp\left(-\frac{e^{rt}-1}{N_0 r}\right) \tag{21}$$

We can then use the binomial probability of the number of pairs *m* (of *k* total allele pairs) with recombination distance greater than the threshold distance to calculate the likelihood of the data given an IBD or recurrent allele.

$$P(m|IBD) = \binom{k}{m} P_{IBD}(D \geq l)^{m}(1-P_{IBD}(D \geq l))^{k-m} \tag{22}$$

$$P(m|rec) = \binom{k}{m} P_{rec}(D \geq l)^{m}(1-P_{rec}(D \geq l))^{k-m} \tag{23}$$

Finally, for each variant, we combine the above likelihoods for the right and left hand sides ($P(m|IBD)^R$ and $P(m|IBD)^L$) to calculate a composite likelihood ratio:

$$\Lambda = log\left(\frac{P(m|IBD)^R P(m|IBD)^L}{P(m|rec)^R P(m|rec)^L}\right) \quad (24)$$

***Application of likelihood-based model simulated data***

We applied the likelihood-based model described above to our simulated IBD and non-IBD variants with uniform recombination rate and no selection. To choose the threshold recombination distance *l*, we calculated the composite likelihood ratio using a range of possible values (*l* = 0.0001, 0.0005, 0.001, 0.011, 0.021, 0.031, 0.041, 0.051, 0.061, 0.071, 0.081, 0.091, 0.02, 0.05, 1.0, 2.0, or 5.0 cM) and chose the distance threshold with the largest AUC to plot in **Supplementary Figure 10**. Chosen values of *l* and AUCs for each allele count are listed in **Supplementary Table 2**.

**Supplementary Table Descriptions**

**Supplementary Table 1.** Areas under curve (AUC) from applying the Bayesian hierarchical model to a range of simulation types. Mean: bootstrapped mean AUC; Median: bootstrapped median AUC; 95CI_L: lower limit of bootstrapped AUC 95% confidence interval; 95CI_U: upper limit of bootstrapped AUC 95% confidence interval; Hotspot_r: simulated hotspot recombination rate (events/bp/generation), $1x10^{-8}$ means there was a uniform recombination rate; AC: allele count; Selection: was selection simulated (logical value).

**Supplementary Table 2.** Areas under curve (AUC) from two alternative methods to identify non-IBD variants, applied to simulated data: runtc, a variant age estimator; and LLR, a likelihood-based approach. Mean: bootstrapped mean AUC; Median: bootstrapped median AUC; 95CI_L: lower limit of bootstrapped AUC 95% confidence interval; 95CI_U: upper limit of bootstrapped AUC 95% confidence interval; AC: allele count; Method: method used in AUC calculation; distance_threshold: for the LLR approach, the distance threshold *l* used to calculate the likelihood.

**Supplementary Table 3.** The correlation between expected and observed fractions of non-IBD variants, for a given allele count and subset of variants. r: Pearson's correlation coefficient; P.value: P-value of r; 95%CI_L: lower limit of 95% confidence interval of r; 95%CI_U: upper limit of 95% confidence interval of r; AC: allele count; Type: variants included (all = all variants, cgt = CpG>T variants only, ncgt = all variants except CpG>T variants).

**Supplementary Table 4.** Expected and observed fractions of variants called non-IBD for 5-mer sequence contexts. Context: sequence context; nVars: number of variants of this allele count and context included; n-nIBD: number of variants called non-IBD; obsFrac: fraction of variants called non-IBD; polyProb: polymorphism probability of this sequence context; expFrac: expected fraction of non-IBD variants of this sequence context; AC: allele count.

**Supplementary Table 5.** Results of linear or logistic regression predicting non-IBD variants from genomic annotations. Estimate: regression beta coefficient estimate; SD: standard deviation of beta estimate; P: P-value of beta estimate; AC: allele count; Method: linear or logistic regression (logistic used non-IBD calls as outcome variable, linear used posterior probability of non-IBD as outcome variable); Input: variants included in model (all or CpG>T only); Annotation: genomic annotation; hsDist.z: hotspot distance z-score; rRate.z: local recombination rate z-score; B.z: McVicker's B statistic z-score; RT.z: replication timing z-score; GC.z: local GC content z-score; mOv.z: ovary CpG methylation z-score; mTes.z: testes CpG methylation z-score; DP.z: read depth z-score; VQSLOD.z: variant quality z-score; polyProb.z: 7-mer polymorphism probability.

**Supplementary Table 6.** Results of logistic regression predicting non-IBD variants in putative gene conversion tracts vs. all other non-IBD variants, with genomic annotations as predictors. Estimate: regression beta coefficient estimate; SD: standard deviation of beta estimate; P: P-value of beta estimate; AC: allele count; Annotation: genomic annotation; hsDist.z: hotspot distance z-score; rRate.z: local recombination rate z-score; B.z: McVicker's B statistic z-score; RT.z: replication timing z-score; GC.z: local GC content z-score; mOv.z: ovary CpG methylation z-score; mTes.z: testes CpG methylation z-score; DP.z: read depth z-score; VQSLOD.z: variant quality z-score; polyProb.z: 7-mer polymorphism probability; posterior: posterior probability of being non-IBD.

**Supplementary Table 7.** Correlation coefficients between posterior probabilities of variants being non-IBD from Bayesian hierarchical model, comparing the model run with non-IBD alpha=20 vs. non-IBD alpha=40. r: Pearson's correlation coefficient; P: correlation coefficient P-value; N: number of variants included; AC: allele count.

### Supplementary Figures

**Supplementary Figure 1. Theoretical distributions of the time to most recent common ancestor.**
The density of TMRCAs for an IBD (green) or recurrent (orange) pair of alleles of allele count 2, 6, or 10; with a sample size of 3,621 and an exponential growth rate of 3%.

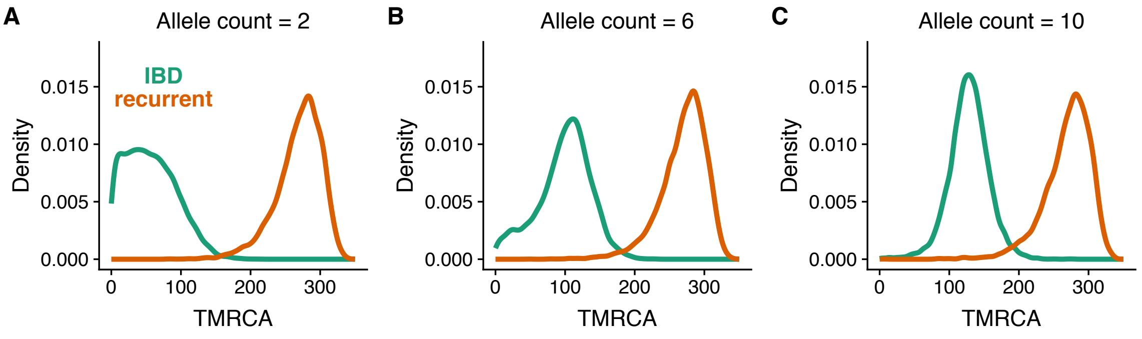


**Supplementary Figure 2. Theoretical distributions of the recombination distance.**
The density of pairwise recombination distances for an IBD (green) or recurrent (orange) pair of alleles of allele count 2, 6, or 10; with a sample size of 3,621 and an exponential growth rate of 3%.

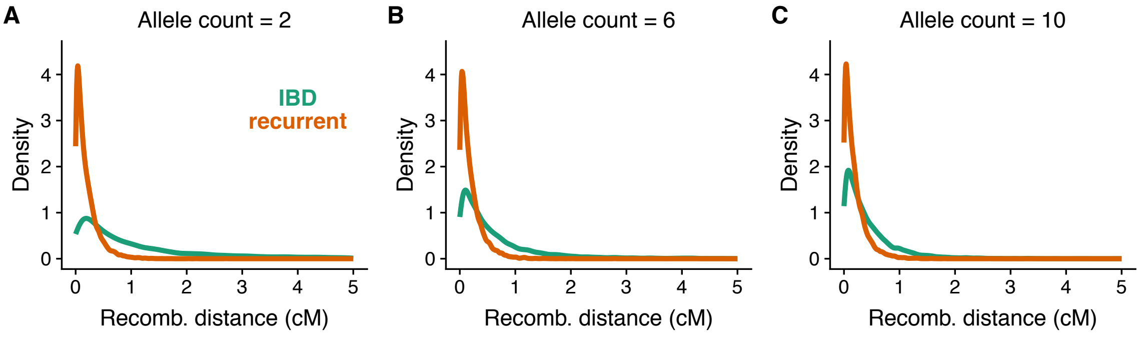

**Supplementary Figure 3. Measuring the obligate recombination distance with unphased diploid genotypes.**
The orange dot represents the focal variant, and blue dots additional variants present in these two individuals. Each individual A and B has two chromosomes (dashed lines), and the red boxes highlight the nearest opposite homozygote genotypes. The purple lines highlight the extent of the shared haplotype of the chromosomes carrying the focal variant. The boxes at the bottom illustrate these two individual's diploid genotypes.

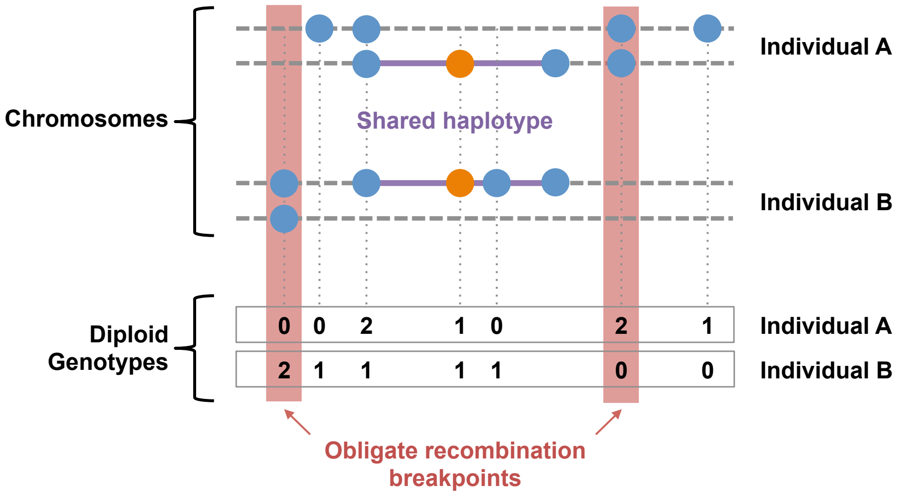


**Supplementary Figure 4. Counts of IBD and recurrent variants in 1000 simulations.**
The counts of IBD and recurrent variants of allele count 2-10 in 1000 SLiM simulations with a uniform recombination rate, uniform mutation rate, and no selection.

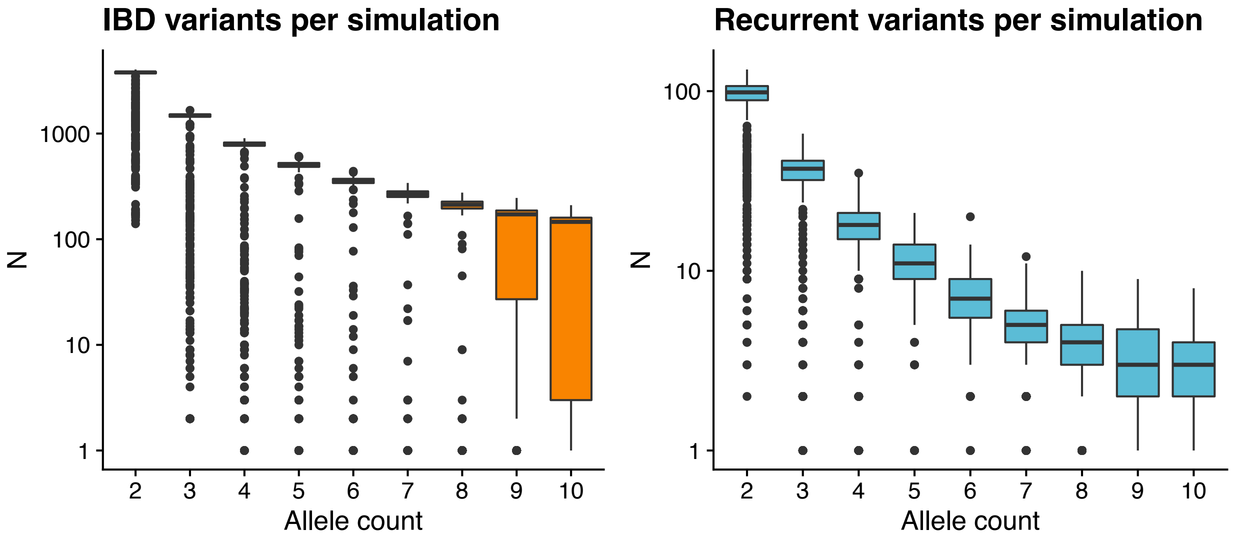

**Supplementary Figure 5. Posterior probabilities of simulated IBD or recurrent variants being non-IBD.**

The empirical cumulative density of posterior probabilities of a variant being non-IBD, for simulated IBD (orange) or recurrent (blue) mutations. Posterior probabilities were generated by application of our Bayesian hierarchical model to variants from simulations with a uniform recombination rate, uniform mutation rate, and no selection.

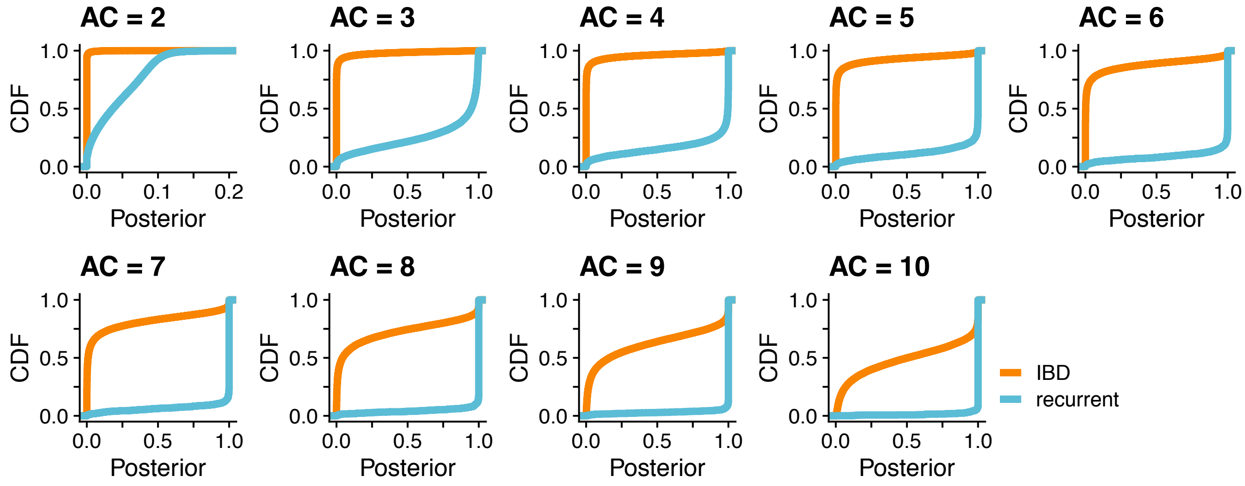


**Supplementary Figure 6. ROC curves of the Bayesian hierarchical model's ability to distinguish IBD and non-IBD variants from simulations with or without selection.**

ROC curves for the Bayesian hierarchical model applied to distinguish simulated recurrent and IBD mutations, with (TRUE) and without (FALSE) the presence of background selection. Each panel represents the application to variants of a given allele count (AC) 2-5. Note the x-axis scale is not the same for AC=2 and the other allele counts.

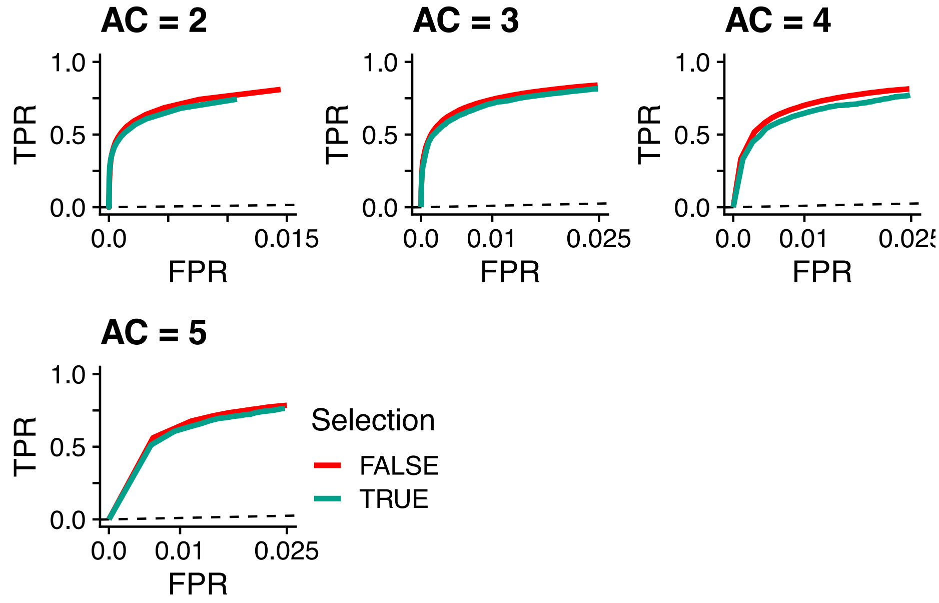

**Supplementary Figure 7. The distribution of allele ages for IBD or recurrent mutations in simulations with and without selection.**

For each allele count, each boxplot describes the distribution of allele ages for simulated IBD (top) or recurrent (bottom) variants, with (TRUE) or without (FALSE) selection present. The recurrent mutation ages are for each independent mutation event, and thus are on average more recent than IBD variants of the same allele count.

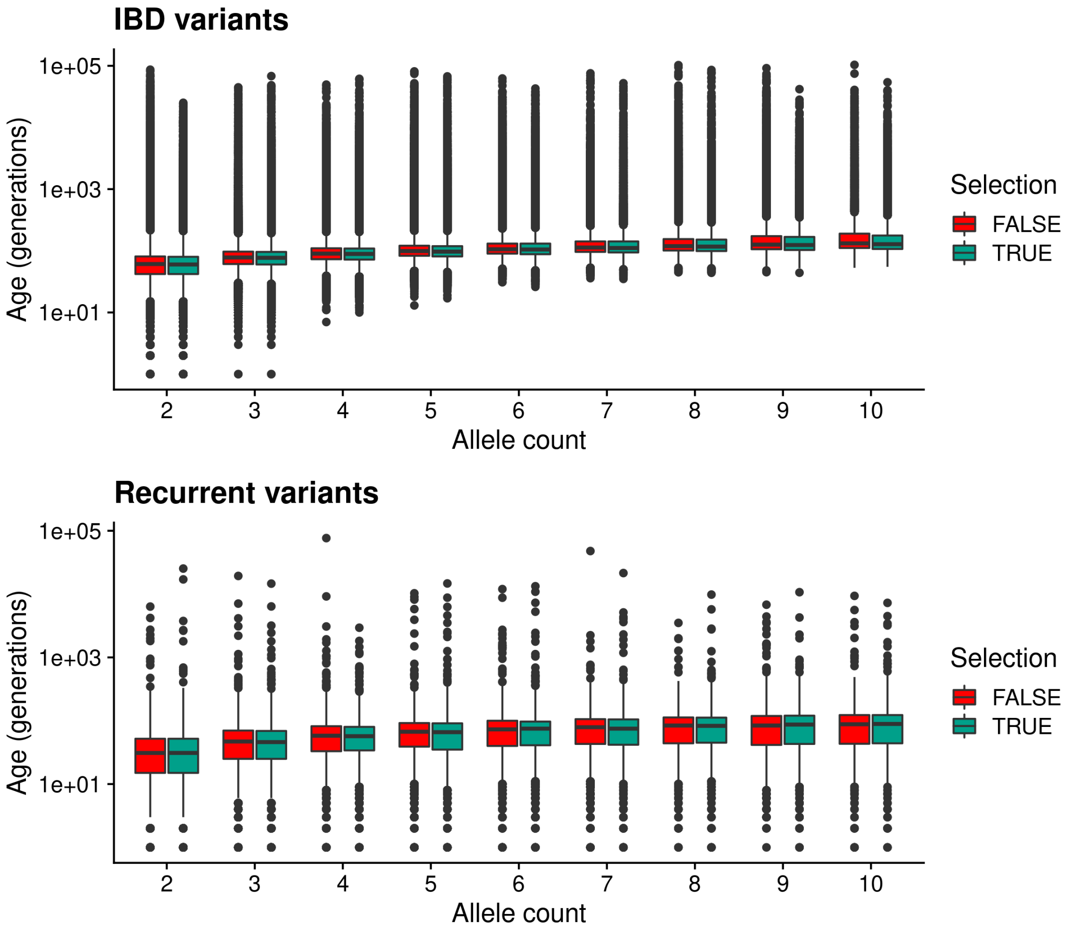

**Supplementary Figure 8. ROC curves of the Bayesian hierarchical model's ability to distinguish IBD and non-IBD variants from simulations with recombination hotspots.**
ROC curves for the Bayesian hierarchical model applied to distinguish simulated recurrent and IBD mutations, for simulations with a range of recombination hotspot strengths. Each panel represents the application to variants of a given allele count (AC) 2-5. Note the x-axis scale is not the same for AC=2 and the other allele counts.

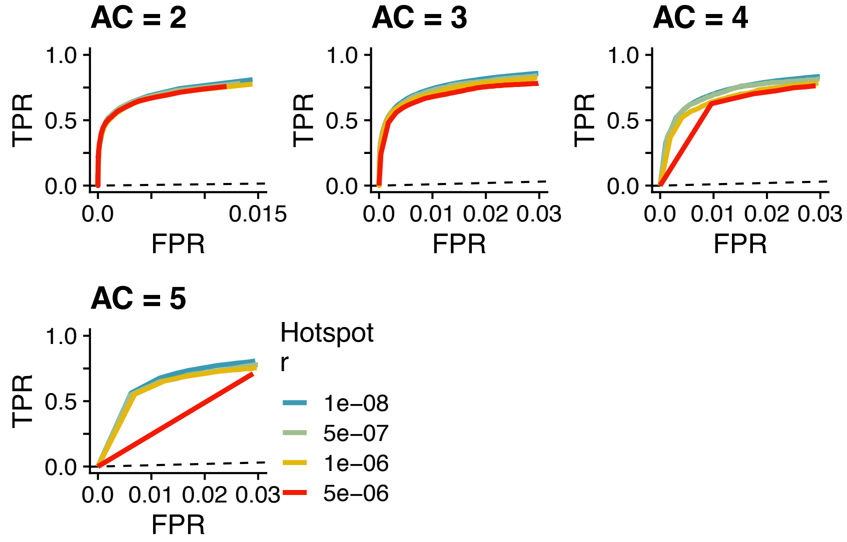


**Supplementary Figure 9. ROC curves comparing the performance of the Bayesian hierarchical model to a theoretical likelihood-based model.**
ROC curves comparing the performance of our Bayesian hierarchical model (BHM; orange) vs. a log-likelihood ratio approach (LLR; blue) to distinguish IBD and recurrent mutations in simulations.

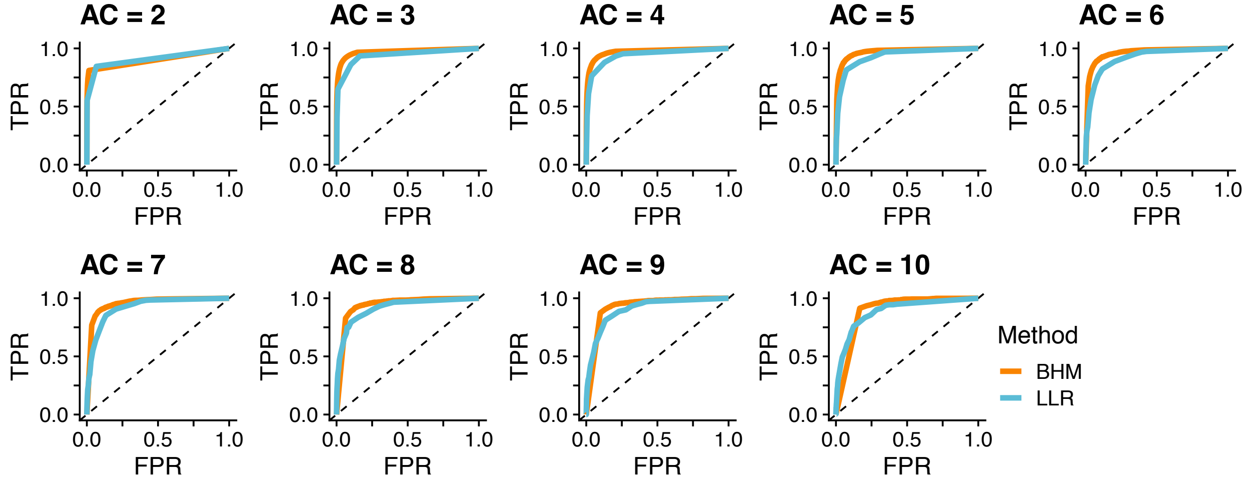

**Supplementary Figure 10. ROC curves comparing the performance of the Bayesian hierarchical model to age estimates from *runtc*.**

ROC curves comparing the performance of our Bayesian hierarchical model (BHM; orange) vs. variant age estimates from *runtc* (blue) to distinguish IBD and recurrent mutations in simulations.

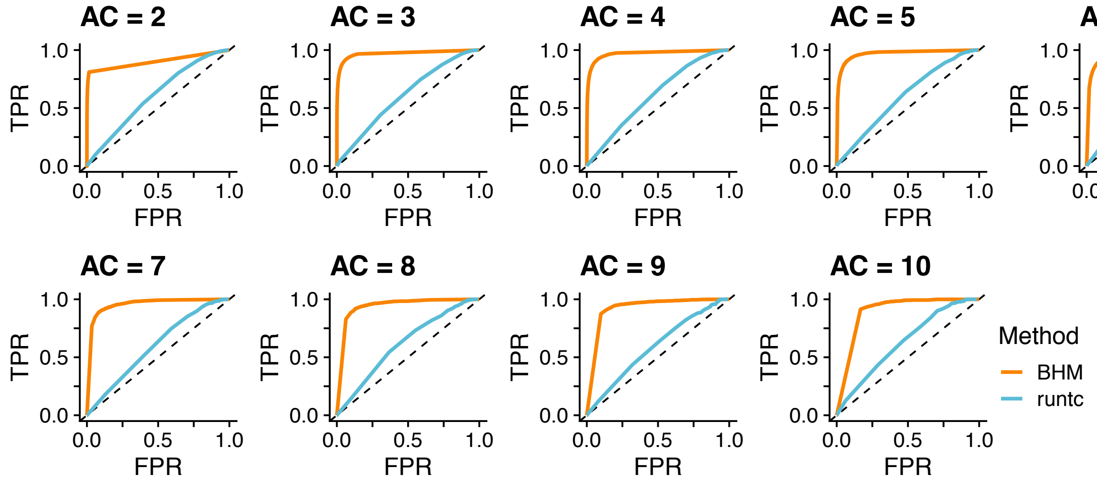


**Supplementary Figure 11. The distribution of posterior probabilities of being non-IBD for biallelic vs. multiallelic variants from UK10K.**

Posterior probabilities were calculated by applying our hierarchical model to biallelic (blue) or multiallelic (orange) sites from the UK10K dataset. Each panel represents variants of a given allele count (AC). CDF: cumulative distribution function.

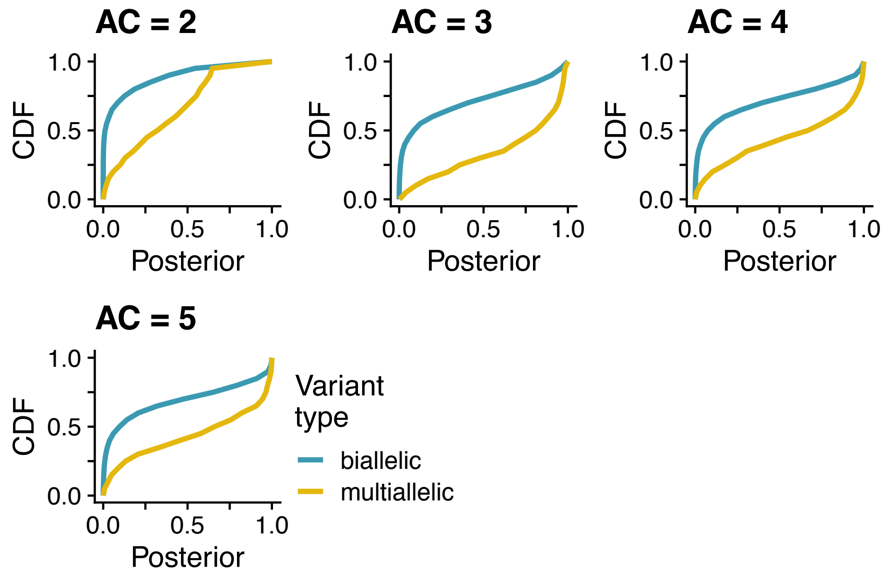

**Supplementary Figure 12. The expected and observed fraction of sites called non-IBD for UK10K variants at CpG sites.**

Each dot represents a 5-mer sequence context. The expected fraction is proportional to the polymorphism probability. Dot colors correspond to 3-mer sequence contexts.

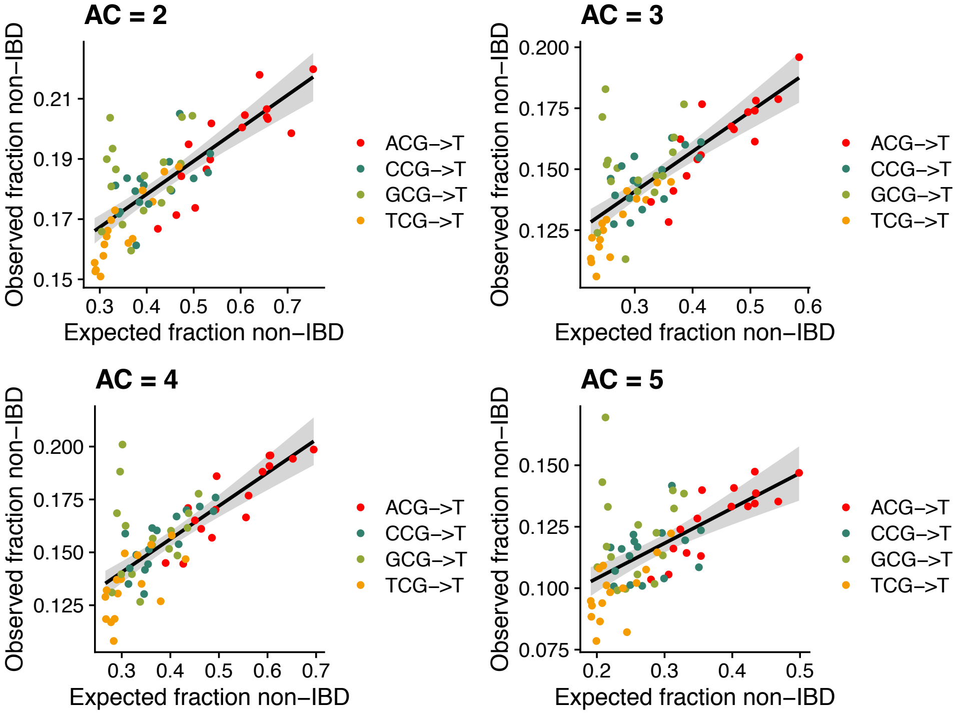

**Supplementary Figure 13. The expected and observed fraction of sites called non-IBD for UK10K variants at non-CpG sites.**

Each dot represents a 5-mer sequence context. The expected fraction is proportional to the polymorphism probability. Dot colors represent the 1-mer sequence context.

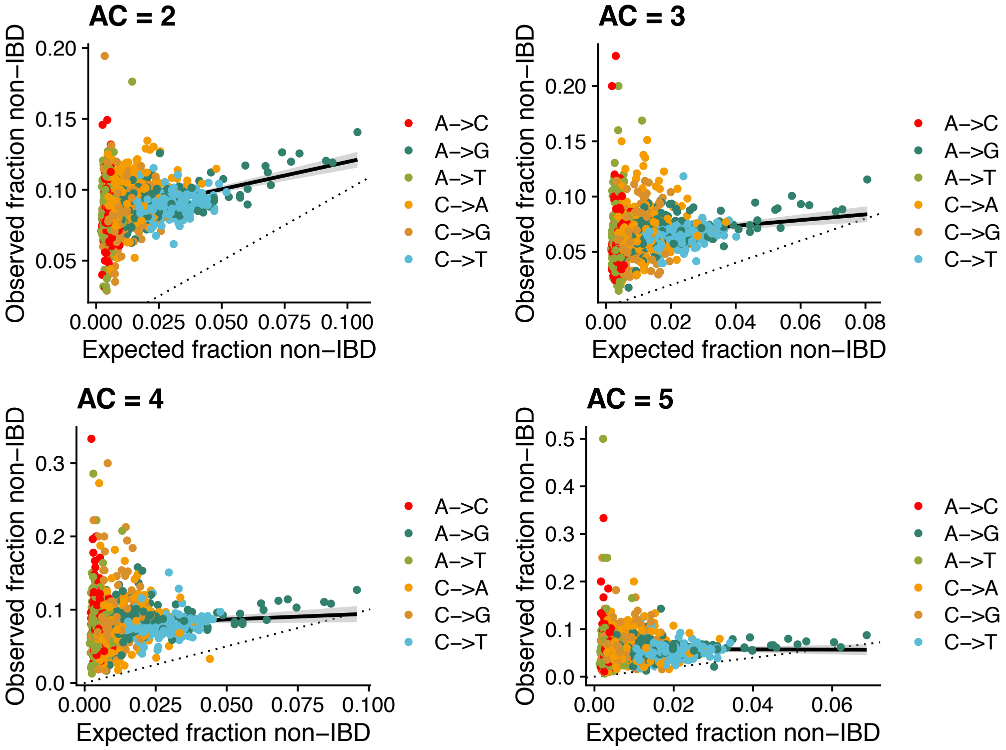

**Supplementary Figure 14. Results of a logistic regression using genomic annotations to predict non-IBD variant calls.**

Logistic regression of genomic annotations (predictor variables) vs. non-IBD variant calls (outcome) for all variant sites. Dot colors represent allele count, and a separate regression was run for variants of each allele count. Each dot's position denotes its beta coefficient estimate, with error bars representing beta ± 1.96*standard error. The vertical dashed line represents a beta estimate of zero. Hotspot distance: physical distance to nearest recombination hotspot z-score; Recombination rate: local recombination rate z-score; B score: McVicker's B statistic z-score; Replication timing: replication timing z-score; GC content: local GC content z-score; Methylation (ovary): ovary CpG methylation z-score; Methylation (testes): testes CpG methylation z-score; Read depth: read depth z-score; VQSLOD: variant quality z-score.

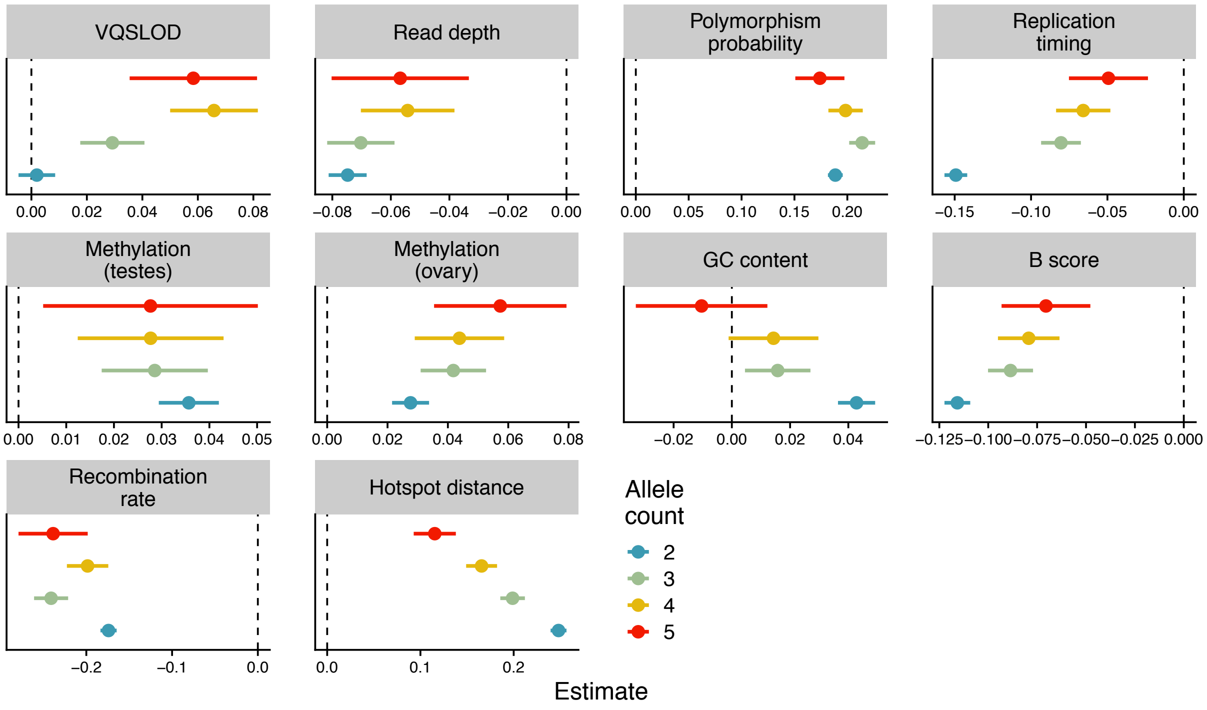

**Supplementary Figure 15. Results of linear or logistic regression using genomic annotations to predict non-IBD variant calls for CpG variants.**

Linear **(A)** or logistic **(B)** regression of genomic annotations (predictor variables) vs. non-IBD variant calls (outcome) for CpG->T sites only, grouped by allele count. Dot colors represent allele count. Each dot's position denotes its beta coefficient estimate, with error bars representing beta ± 1.96*standard error. The vertical dashed line represents a beta estimate of zero. Hotspot distance: physical distance to nearest recombination hotspot z-score; Recombination rate: local recombination rate z-score; B score: McVicker's B statistic z-score; Replication timing: replication timing z-score; GC content: local GC content z-score; Methylation (ovary): ovary CpG methylation z-score; Methylation (testes): testes CpG methylation z-score; Read depth: read depth z-score; VQSLOD: variant quality z-score.

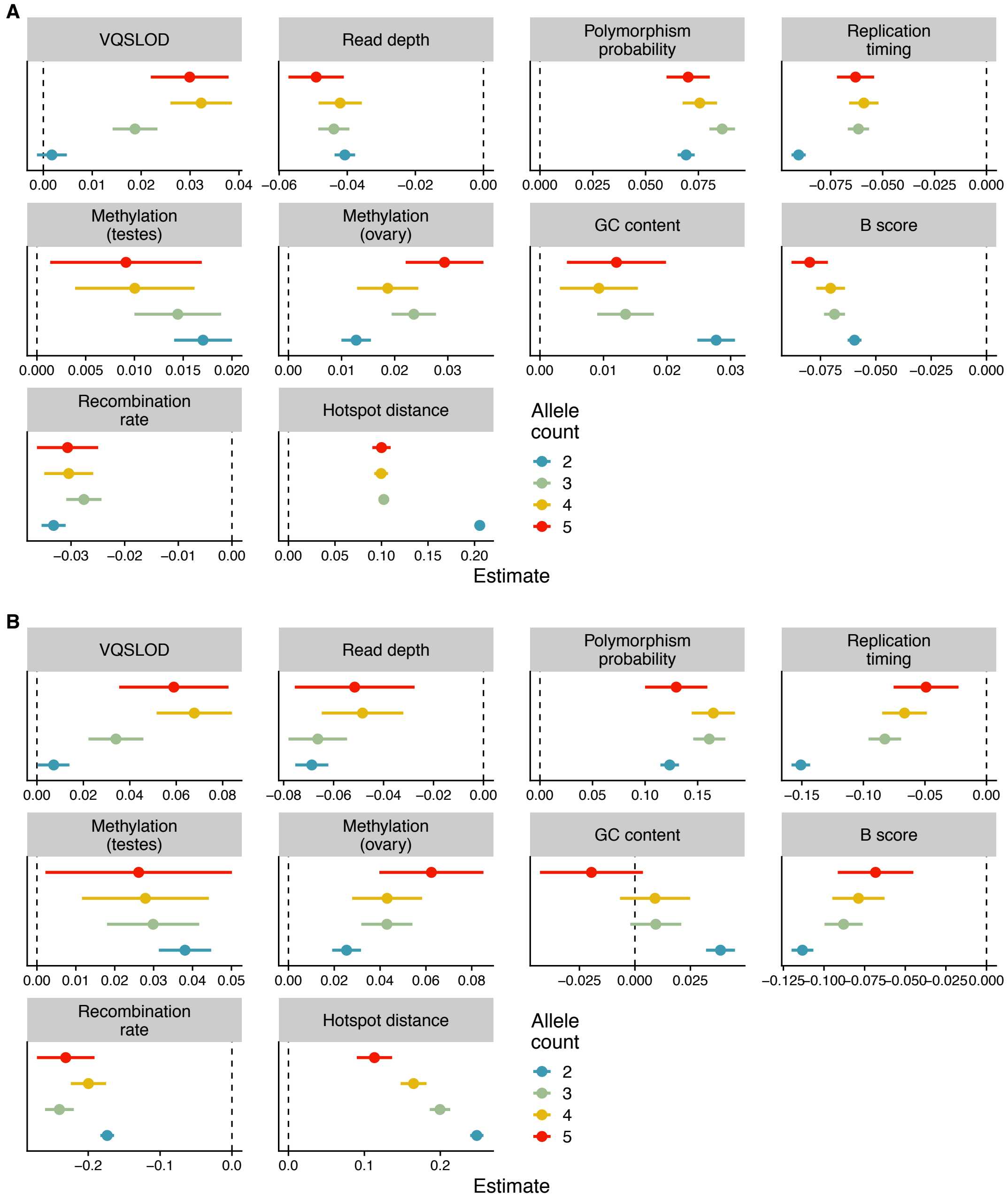

**Supplementary Figure 16. Distribution of putative gene conversion tract lengths.**
Each box plot represents the distribution of tract lengths for putative gene conversion tracts for a given allele count and number of variants in the tract. Note that our heuristic approach to identify putative gene conversions required a maximum length of 1000 base pairs.

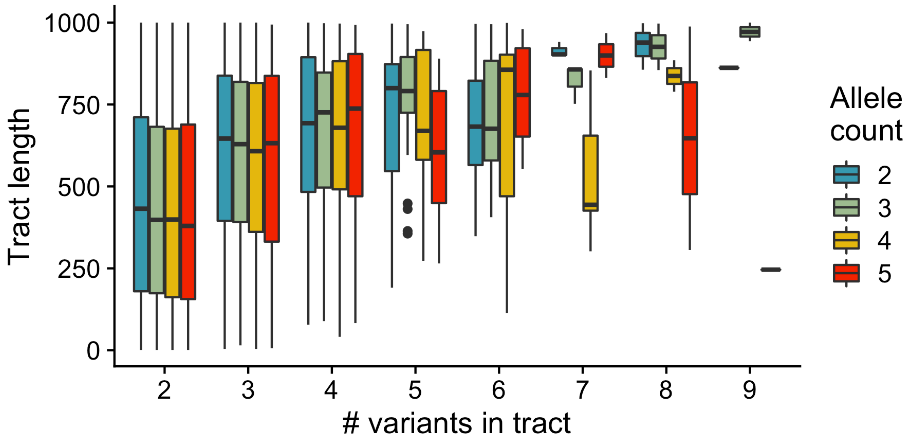
